# Directed differentiation of EA/TEF patient-derived induced pluripotent stem cells into esophageal epithelial organoids reveal SOX2 dysregulation at the anterior foregut stage

**DOI:** 10.1101/2022.02.14.480418

**Authors:** Suleen Raad, Anu David, Melanie Sagniez, Zakaria Orfi, Nicolas A. Dumont, Martin Smith, Christophe Faure

## Abstract

A series of well-regulated cellular and molecular events result in the compartmentalization of the anterior foregut into the esophagus and trachea. Disruption of the compartmentalization process leads to esophageal atresia/tracheoesophageal fistula (EA/TEF). Therefore, the objective is to differentiate pluripotent stem cells (PSCs), namely, embryonic stem cells and iPSCs from healthy individuals and iPSCs from EA/TEF type C patients, into mature 3-dimensional esophageal organoids expressing Involucrin, Keratin-4, -13, and p63. CXCR4, SOX17, and GATA4 expression was similar in both patient and healthy endodermal cells. Key transcription factor SOX2 was significantly lower in patient-derived anterior foregut. RNA sequencing revealed critical genes GSTM1 and RAB37 to be significantly lower in patient-derived anterior foregut. Furthermore, we observed an abnormal expression of NKX2.1 in the patient-derived mature esophageal organoids. We therefore hypothesize that a transient dysregulation of SOX2 and the abnormal expression of NKX2.1 in patient-derived cells could be responsible for the abnormal foregut compartmentalization.

## Introduction

The esophagus and trachea originate from the endodermal diverticulum in the anterior foregut tube. Well-regulated and organized cellular and molecular events result in the separation of the anterior foregut tube into the esophagus and trachea (Raad et al. 2020) (Billmyre, Hutson, and Klingensmith 2015). Disruption of the compartmentalization process results in severe esophageal congenital anomalies such as esophageal atresia with or without tracheoesophageal fistula (EA/TEF), affecting 1 in 3,000 newborns (van Lennep et al. 2019). Several types of EA/TEF have been described based on the location of the malformation and the affected structures, with the most common being type C (>80% of cases), where the upper segment of the esophagus ends in a blind pouch, and a fistula connects the lower part to the trachea. Other less common subtypes include type A (8-10% of cases) where no fistula exists, but the esophagus is disconnected (Clark 1999). EA/TEF-associated anomalies (cardiac, anal, renal, limb, or vertebral) are also reported in 30-50% of syndromic cases. Monogenetic causes account for a minority of EA/TEF cases (<5%) most often in syndromic cases such as Anophthalmia-Esophageal-Genital (AEG) syndrome (*SOX2* mutations), Feingold syndrome (*MYCN* mutations), CHARGE syndrome (*CHD7* mutations), Pallister-Hall syndrome (*GLI3* mutations) and Mandibulofacial dysostosis (*EFTUD2* mutations)(Stoll et al. 2009). Studies have also shown a multigenic architecture of rare variants in several genes, which discriminate EA/TEF cases from controls (Wang et al. 2021). However, the cause of EA/TEF remains largely unknown, and rare genetic variants are seldom reported in non-syndromic, isolated cases. EA/TEF is thus considered a multifactorial anomaly resulting from genetic and environmental factors (Brosens et al. 2014).

During embryogenesis, the esophagus and trachea arise after the separation of the anterior foregut endoderm common tube at week 4-5 in humans and embryonic day 9.5-11.5 in mice. In animal models (mouse and Xenopus), the dorsal/ventral patterning of the anterior foregut allows spatial specification of the two presumptive organs: the esophagus on the dorsal side of the anterior foregut tube (characterized by the expression of the transcription factor SOX2) and the trachea on the ventral side of the foregut tube (characterized by the expression of the NKX2.1 gene) (Minoo et al. 1999; Que et al. 2007; Kim et al. 2019). Studies in mice have also demonstrated that the dorsal/ventral patterning is initiated by gradual expression of mesodermal Wnt2/2b, Bmp4, and Noggin along the dorsal-ventral axis. Bmp signaling pathway inhibit SOX2 expression on the dorsal side of the anterior foregut (Domyan et al. 2011) and drive Nkx2.1 expression toward the tracheal lineage. Functional genomic studies in mice and Xenopus have also been used to mimic the genetics and morphogenetic regulation of normal and abnormal foregut compartmentalization representing human esophageal anomalies such as the EA/TEF (Kim et al. 2019; Fausett and Klingensmith 2012; Raad et al. 2020). However, these studies utilize methods resulting in functional loss of specific genes that may not represent the genetic complexity observed in humans. The esophagus of the human differs structurally and morphologically from the esophagus of the mouse (Rosekrans et al. 2015). Therefore, there is a need to have a representative model of human esophagus development to not only decipher but also understand the possible mechanisms leading to EA/TEF. Induced pluripotent stem cells (iPSCs) offer an excellent tool in gaining insights not only on human embryonic and developmental ontologies but also to model diseases (Karagiannis et al. 2019; Rowe and Daley 2019) through the directed differentiation to specific organs originating from all three germ layers. To date, patient-derived iPSCs have not yet been used to study digestive malformations. Recently, patient-derived iPSCs have been used to study congenital heart diseases, where intrinsic defects were observed in differentiated cardiomyocytes derived from these iPSCs (Miao et al. 2020; Hrstka et al. 2017; Yang et al. 2017). Over the last few years, studies using healthy human iPSCs have been used to generate mature esophageal epithelium (Zhang et al. 2018; Trisno et al. 2018) and confirmed previous findings that SOX2 is key to promoting esophageal specification and the critical role of BMP, TGF-ß, and WNT signaling pathways during esophageal development (Que et al. 2007; Teramoto et al. 2020; Zhang et al. 2018; Trisno et al. 2018; Que et al. 2006; Li et al. 2007; Woo et al. 2011; Domyan et al. 2011).

Therefore, the objective of the current study is to differentiate embryonic stem cells (ESCs) and iPSCs from healthy individuals and iPSCs from three isolated EA/TEF type C pediatric patients into mature esophageal organoids in matrix- and xenogeneic-free culture conditions. We adapted and modified a stepwise differentiation protocol (Matsuno et al. 2016; Zhang et al. 2018; Giroux et al. 2017; DeWard, Cramer, and Lagasse 2014) by manipulating key signaling pathways involved in esophagus development. We investigated the gene and protein expression profiles of key signaling molecules in patient cells and compared them to healthy cells at each developmental stage. Furthermore, by combining targeted gene expression and nanopore RNA sequencing, we demonstrated that patient-derived cells exhibit unique molecular signatures, especially at the anterior foregut stage. Our study establishes a basic framework to understand the morphogenesis and mechanisms involved during early esophageal development by using patient-derived iPSCs.

## Results

Our modified protocol includes the stepwise differentiation of PSCs into mature esophageal organoids with checkpoints at four developmental stages: (i) definitive endoderm, (ii) anterior foregut, (iii) mature esophageal epithelium and (iv) 3-dimensional esophageal organoids **(Scheme 1).** We compared the differentiation potential to these stages between healthy PSCs (embryonic stem cell line [H9] (female)) and an iPSC derived from a non-familial healthy male and EA/TEF iPSCs (2 males and 1 female).

**Scheme 1:**
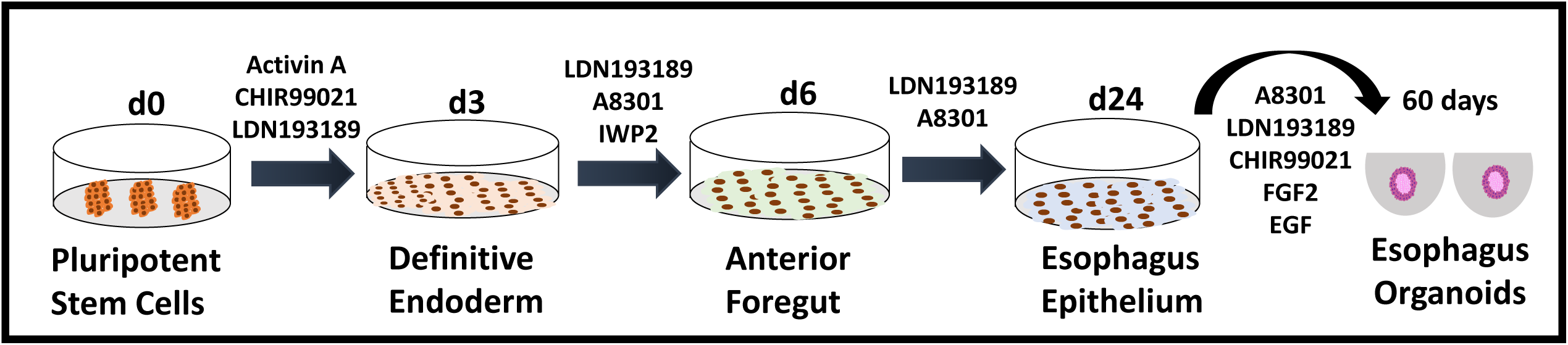
Stepwise differentiation protocol of human pluripotent stem cells into esophagus organoids. Illustration of different signaling pathways manipulated to differentiate pluripotent stem cells into each developmental stage starting from definitive endoderm, anterior foregut, esophagus epithelium and esophagus organoids. **Activin A:** dimeric growth and differentiation factor activates Nodal/TGFb pathway **CHIR99021:** aminopyrimidine derivative that is potent to GSK3 inhibitor, a key inhibitor of the WNT pathway. CHIR99021 activates the WNT pathway **LDN193189:** dihydrochloride potent and selective ALK2 and ALK3 inhibitor. It inhibits BMP4 mediated Smad1/5/8 activation **A8301:** potent inhibitor of the TGFb type I receptor ALK5 kinase, type I activin/nodal receptor ALK4 and type I nodal receptor ALK7 **IWP2:** inhibits WNT pathway at the level of the pathway activator Porcupine, leading to WNT secretion and signaling capability **FGF2:** growth factor belonging to the FGF superfamily. It stimulates cell proliferation. **EGF:** potent growth factor belonging to the EGF family. It induces cell proliferation, differentiation, and survival.

### 1. Derivation of EA/TEF patient iPSCs

iPSC lines from 3 pediatric isolated type C EA/TEF patients without any associated malformations were established by reprogramming peripheral blood mononuclear cells (PBMCs) in the Stem Cell core facility at CHU-Sainte Justine. Their pluripotency was confirmed by the mRNA expression of pluripotent genes SOX2, NANOG, and OCT4 and immunofluorescence staining with SOX2, NANOG, OCT4, and SSEA4. All 3 iPSC cell lines had a normal karyotype, had no pathogenic genetic variants in established EA/TEF risk genes and showed the ability to differentiate into the three germ layers as evidenced through teratoma formation. **(Raad et al. 2021, submitted to Stem Cell Research).**

### 2. Similar differentiation potential of healthy and EA/TEF patient PSCs into definitive endodermal (DE) cells

The first critical step is the differentiation of PSCs into endodermal cells that give rise to the entire epithelial lining of the gastrointestinal tract, including the esophagus epithelium(Wells and Melton 1999). We evaluated the efficiency of endoderm differentiation by qPCR through gene expression of specific markers CXCR4, GATA4, and SOX17 (**Fig. 1A-D**). There was no significant difference in gene expression levels between healthy and patient-derived definitive endoderm. At the protein level, CXCR4 and GATA4 were observed in the cytoplasm, whereas SOX17 was observed in the cell nuclei, confirming DE commitment in both groups (**Fig. 1E**). Furthermore, we also verified that no ectodermal cells were present through the absence of OTX2 **(Data not shown).**

**Figure 1:**
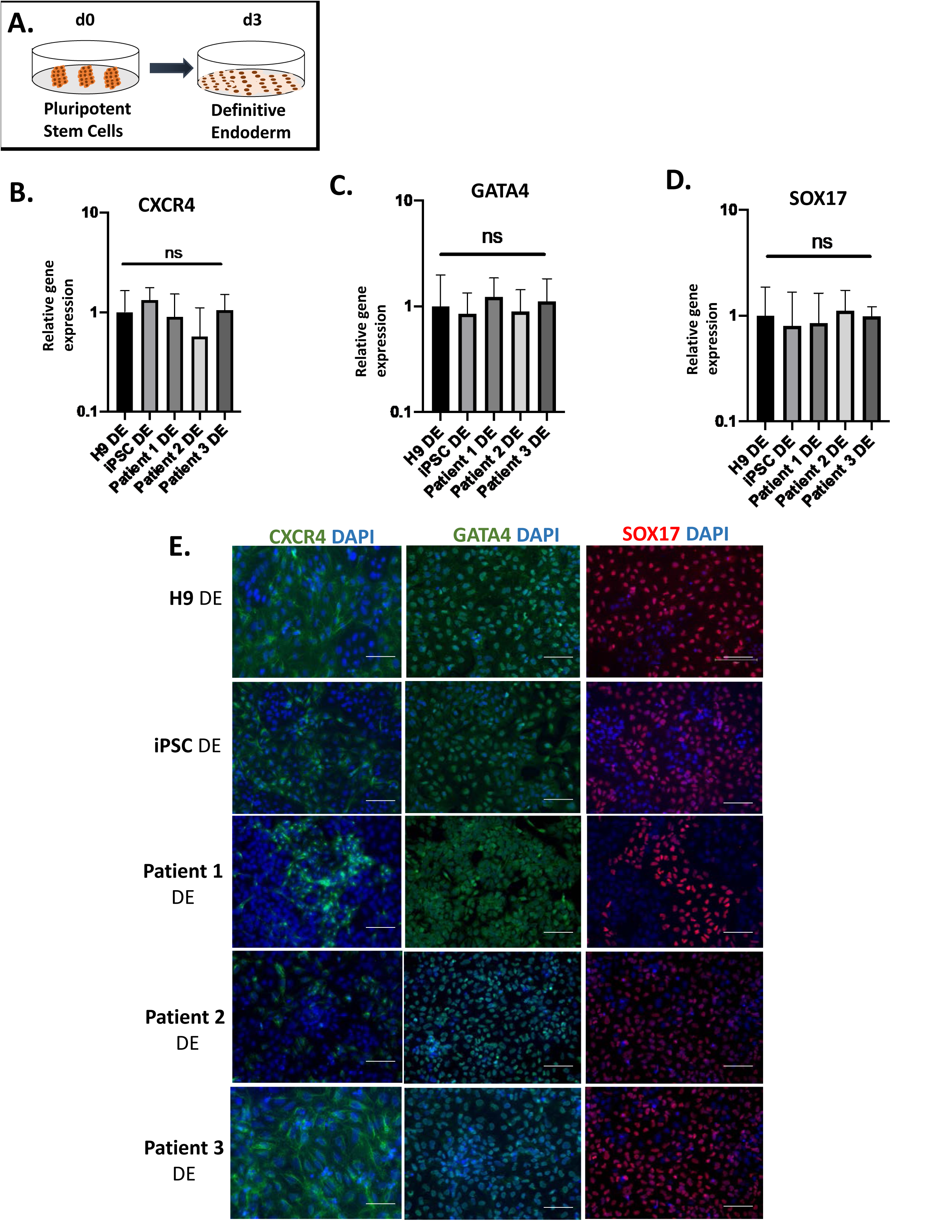
Differentiation of healthy and EA/TEF patient-derived pluripotent stem cells into definitive endoderm (DE). **A)** Illustration of first stage of differentiation into definitive endoderm. **B, C and D)** Expression of key endodermal markers CXCR4, GATA4 and SOX17 was quantified by qPCR and reported as fold change. The fold change was generated by normalizing the transcript levels to those of healthy (H9) DE cells. Data representative mean ±SEM (n>3 technical replicates for each biological cell line). **E)** DE cells from healthy and patients express similarly CXCR4, GATA4 and SOX17 at the protein level by immunofluorescence staining. Negative controls were included in each staining. Scale bar 50um.

### 3. Critical dorsal esophageal marker is downregulated in EA/TEF patient-derived anterior foregut cells

Developmentally, the anterior side of the foregut tube separates dorsally into the esophagus and ventrally into the trachea and therefore, to generate the dorsal side of the anterior foregut, we inhibited key signaling pathways shown to be critical for esophagus specification; the BMP, TGFb, and WNT pathways (**Fig. 2A**). Anterior foregut cells in both groups expressed PAX9, a foregut endodermal marker at the gene and protein levels (**Fig. 2B, D**). The cells also expressed ISL1 (**Fig. 2C, E**), a recently identified critical marker that contributes to the specification of anterior foregut to both esophageal and tracheal epithelium (Kim et al. 2019). ISL1 regulates the expression of NKX2.1 and is required for esophagus-trachea separation (Zhang et al. 2018; Kim et al. 2019). The expression profiles of ISL1 and PAX9 were similar in both groups (**Fig. 2D, E**).

**Figure 2:**
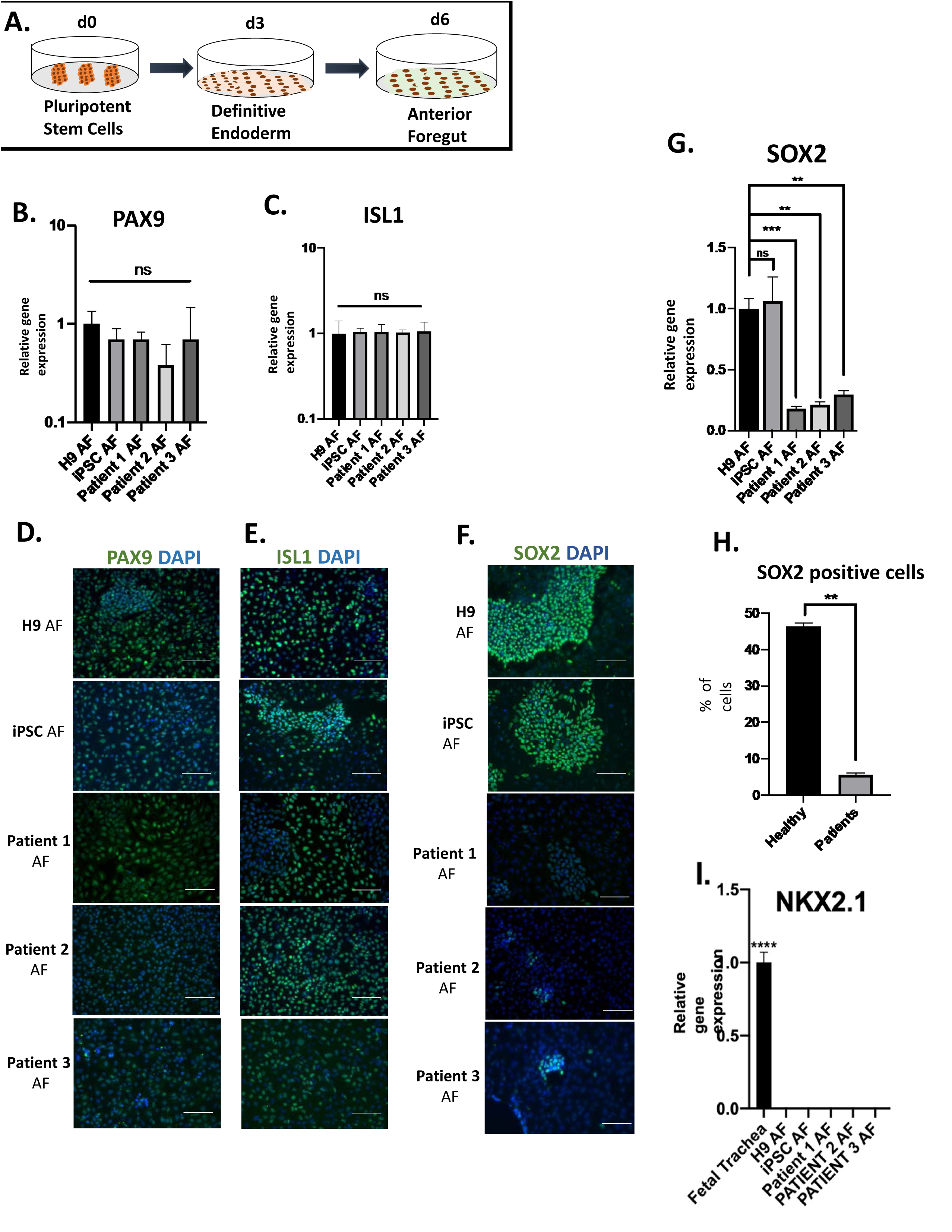
SOX2 expression significantly downregulated in patient-derived Anterior foregut (AF) cells. **A)** Schematic representation of differentiation from definitive endoderm into anterior foregut. **B and C)** PAX9 and ISL1 are expressed similarly by qPCR in both healthy and patient derived anterior foregut cells. The fold change was generated by normalizing the transcript levels to those of healthy (H9) DE cells. Data representative mean ±SEM (n>3 technical replicates for each biological cell line). **D and E)** Healthy and patient-derived anterior foregut cells express PAX9 and ISL1 nuclear staining by immunofluorescence similarly. **F)** Low expression of SOX2 at the protein level by immunofluorescence in all 3 patients. To note, patients 2 and 3 have fewer cells with bright fluorescence and patient 1 has more cells with less fluorescence **G)** SOX2 is downregulated in all three patient-derived anterior foregut cells. Transcript levels were compared to those of H9 AF. Data represents mean ± SEM (n>3 technical replicates for each biological cell line) **p<0.01, *** p<0.001 by unpaired two tailed student’s t test **H)** Cell counting of positive cells using Image J software to count positively stained cells revealed that less than 10% of the patient AF cells are positive for SOX2. Scale bar 50um. **I)** Lack of expression of NKX2.1 at the AF stage in both groups. The fold change was generated by normalizing the transcript levels to those of healthy (H9) DE cells. Data representative mean ±SEM (n>3 technical replicates for each biological cell line)

However, SOX2, a critical transcription factor necessary for foregut morphogenesis and expressed on the dorsal side of the anterior foregut, was downregulated in the patient-derived cells (**Fig. 2F, G**). Following quantification, we observed that SOX2 expression was significantly lower in all three patients (∼10%) compared to the healthy foregut cells (45%) (**Fig. 2F, H**). It is known that disruption of SOX2 expression leads to an abnormal separation of the anterior foregut into the esophagus and trachea (Teramoto et al. 2020).

At the anterior foregut stage when the compartmentalization occurs, two critical transcription factors namely, SOX2 and NKX2.1, have a reciprocal repressive function. Nkx2-1 binds to silencer sequences near the SOX2 gene and represses its transcription (Kuwahara et al. 2020; Kim et al. 2019; Trisno et al. 2018; Han et al. 2020). However, dysregulated SOX2 did not affect the expression of NKX2.1, which remained undetected in both groups (**Fig. 2I**). Furthermore, at this stage, we also confirmed the absence of other lineage markers, specifically, a mid-hindgut marker, CDX2 **(Data now shown)** and a posterior foregut marker, HNF4a **(Supplementary Fig. S1A)**

### 4. Novel transcript isoforms and distinct molecular signatures in patient-derived anterior foregut cells using nanopore sequencing

Low SOX2 expression levels in patient-derived anterior foregut cells led us to investigate what other genes could be involved in this stage of development. We thus applied RNA sequencing (Oxford Nanopore) on the anterior foregut cells derived from the two groups to quantify gene expression globally and characterize RNA isoform diversity across the surveyed samples. We report 2 new isoforms of the SOX2 gene, one of which presents an intron in its 3’UTR. This filtered *de novo* assembly was used as a reference for sample-specific abundance estimation. The latter revealed a potential batch-effect associated with 2 separate sample preparations that, unfortunately, also coincided with the sex of the individual, representing 42% of the observed variation in the data **(PC1,** **Fig. 3A**). We performed batch correction using SVA (Leek et al. 2021) to mitigate this effect, which resulted in an effective separation of disease and healthy samples across the 2 remaining principal components (**Fig. 3B**). Sequencing validated our previous observations that SOX2 is less expressed at the anterior foregut stage in the patient group. The differential transcript expression with DeSeq2 identified 173 transcripts that presented an over 2-fold change in normalized expression with a p-value below 0.01 **(Supplementary Table 1).** We could identify gene expression signatures unique to both conditions (**Fig. 3C, D**). Specifically, both SOX2 transcripts that overlap the TaqMan probes used in qPCR (SOX2-201 and SOX2-201(o)-25276.2, **Fig. 3D**) presented an average TPM of 618 in healthy samples versus 253 in affected samples. GSTM1-201 was among the top differentially expressed isoforms in patient-derived cells (log2(fold change) = 5.61). It is significantly lower in all 3 patients compared to healthy samples (**Fig. 3D**). Previous work has shown that GSTM1 is associated with EA/TEF (Filonzi et al. 2010). Also, among the list is RAB37-204 (log2(fold change) = 1.08), an endosomal protein critical for vesicle trafficking regulation. Rab proteins have been previously linked to foregut malformations (Nasr et al. 2019; Nam et al. 2010; Edwards et al. 2021). Among the top differentially expressed transcripts are several non-coding RNAs, including Y-RNA, MEG3 (**Fig. 3D** **and Supplementary Table 1).**

**Figure 3:**
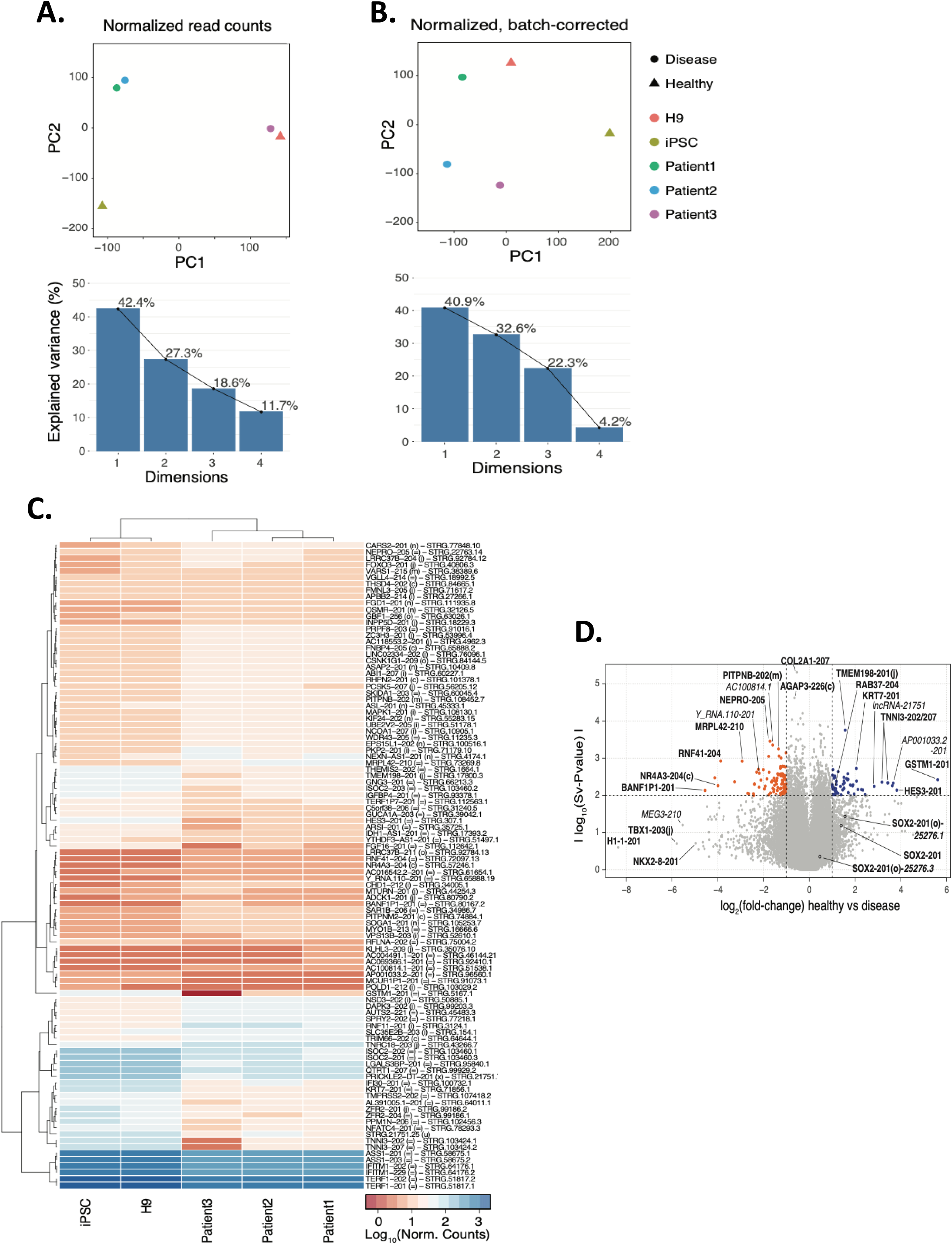
De novo assembly from long read RNA sequencing of patient versus healthy cells. **A)** Principal component analysis of sequencing data before and after **(B)** replicate- and sex-specific batch correction. **C)** Heatmap of the 100 highest fold-change transcripts (batch-corrected P-value< 0.01). **D)** Volcano-plot of differentially expressed transcripts (horizontal threshold set at batch-corrected P-value = 0.01 and vertical thresholds set at log2 (FoldChange) ±1).

### 5. NKX2.1, a tracheal marker, is expressed in EA/TEF patient-derived esophageal epithelium

We further differentiated the anterior foregut cells into esophagus epithelium by inhibiting the BMP and TGF-ß pathways (Que et al. 2006; Guyot and Maguer-Satta 2016) (**Fig. 4A**). Even with low SOX2 expression in patient-derived anterior foregut cells, we observed that these cells were committed to an esophageal fate. Specifically, we observed that esophageal epithelial cells derived from both groups expressed esophageal marker, p63, normally expressed in the basal proliferative layer of the developing esophagus (**Fig. 4B**). Cells from both groups also expressed KRT4, an esophageal squamous epithelial marker (**Fig. 4C, E**). Interestingly, SOX2, a marker also expressed by the basal proliferative esophageal epithelium, was observed at similar levels in both groups (**Fig. 4D, E**).

**Figure 4:**
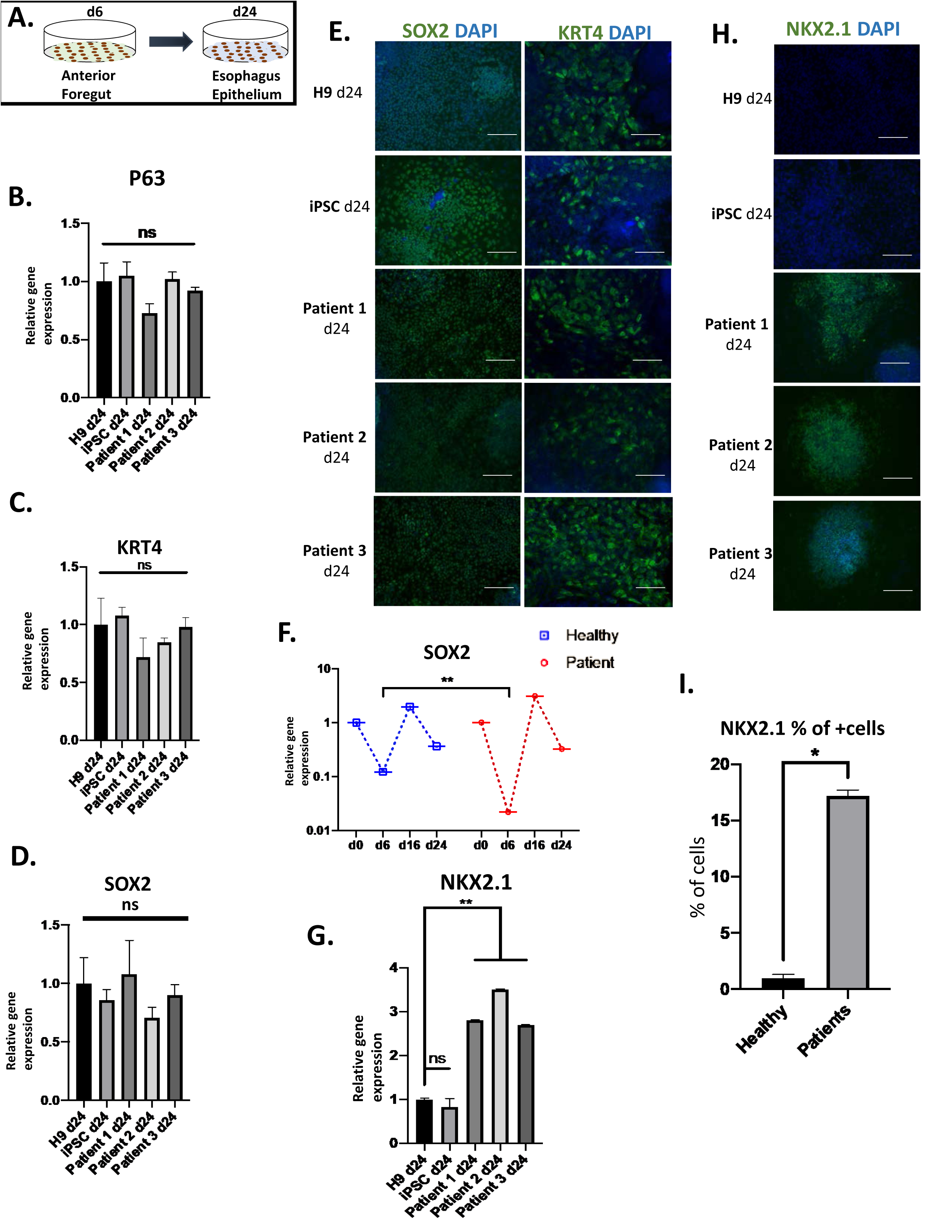
Esophagus epithelial cells derived from EA/TEF patients express NKX2.1, a tracheal marker. **A)** Illustration representing the differentiation of anterior foregut cells into esophagus epithelium **B, C and D)** Transcript levels of p63, KRT4 and SOX2 shows similar expression in both healthy and patient groups. Relative expression was compared to that of H9 d24 esophagus epithelium. The fold change was generated by normalizing the transcript levels to those of healthy (H9) DE cells. Data representative mean ±SEM (n>3 technical replicates for each biological cell line). **E)** Immunofluorescence staining of d24 esophagus epithelium confirming the expression of SOX2 and KRT4 similarly in healthy and patient derived esophagus epithelium **F)** Relative expression of SOX2 throughout differentiation. The fold change was generated by normalizing the transcript levels to those of healthy (H9) DE cells. Data representative mean ±SEM (n>3 technical replicates for each biological cell line) **p<0.01, *** p<0.001 by unpaired two tailed student’s t test. **G)** Abnormal expression of NKX2.1 in patient derived esophagus epithelium. Transcript levels were significantly higher in all 3 patient cells. Transcript levels were compared to that of H9 esophagus epithelium Data represents mean ±SEM (n>3 technical replicates for each biological cell line) **p<0.01, *** p<0.001 by unpaired two tailed Student’s t test. **H)** At the protein level, NKX2.1 was also expressed in patient-derived esophagus epithelial cells, confirming our findings at the RNA level. **I)** Cell counting of NKX2.1 positive cells by the software Image J counting cells positively staining for NKX2.1. Scale bar 50um

The expression of SOX2 during esophagus differentiation of EA/TEF patient iPSCs differs greatly from healthy group. At the anterior foregut stage, we observed a temporal downregulation of SOX2 expression in the 2 groups but was significantly more pronounced in patient cells (**Fig. 4F**). Interestingly, SOX2 expression goes back to similar levels to that of the healthy group at the esophageal epithelial stage (**Fig. 4F**). However, at this stage though the SOX2 expression returned to normal levels we observed a significantly higher expression of NKX2.1 in patient-derived esophageal epithelial cells at the gene and protein levels (**Fig. 4G, H**). 17% of the cells were positive for NKX2.1 in patient-derived esophageal epithelium (**Fig. 4I**). A recent study (Kim et al. 2019) identified ISL1 to be a regulator of NKX2.1 during foregut separation. But we did not observe any significant difference in the expression of ISL1 in both healthy and patient derived cells at both the anterior foregut and esophageal epithelial stage **(Supplementary Fig. S1B).**

### 6. Mature esophageal epithelial organoids express key markers: Involucrin (INV), Keratin-4 and -13 (KRT-4 and KRT-13), and p63

For further maturation and to allow for cellular organization of the esophageal epithelium into a stratified squamous epithelium, 3-dimensional organoids were generated and further matured. Cells were detached from their 2-dimensional culture conditions, matured in suspension (**Fig. 5A**) and within 48 hours the cells clustered together to form organoids (**Fig. 5B**). We observed no morphological and proliferative differences between the healthy and patient-derived organoids **(Fig. 5B, Supplementary Fig. S2).** We observed high gene and protein expression of suprabasal markers such as KRT4, KRT13, and Involucrin in both healthy and patient-derived organoids closer to the fetal esophagus than to adult esophagus biopsy (**Fig. 5C, D, E, F**) in addition to p63, a basal proliferative marker (**Fig. 5G, H**). We expected the expression of these markers to be normal at this stage because the upper and lower end of the esophagus in EA/TEF patients is not affected morphologically.

**Figure 5:**
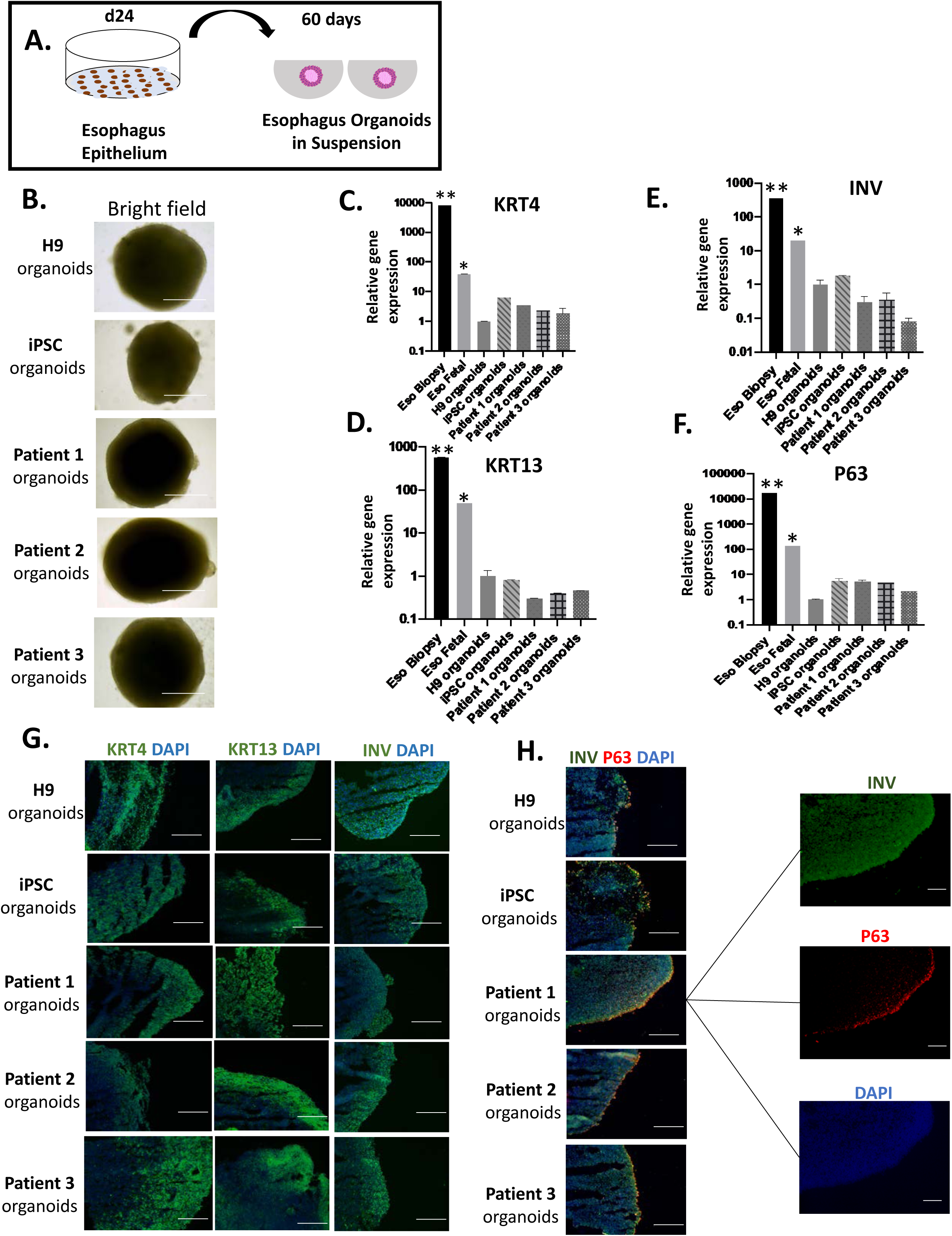
Healthy and EA/TEF patient PSCs can generate mature esophagus organoids. **A)** Illustration of differentiation from esophagus epithelium progenitors into esophagus organoids. **B)** Bright field images of healthy and patient-derived esophagus organoids reveal similar morphology between the two groups. **C, D, E, F)** transcript levels of mature esophagus markers such as KRT4, KRT13, INV and p63 reveal a similar expression between healthy and patient-derived esophagus organoids. Relative expression was compared to H9 derived esophagus organoids. Two references were included in the graphs, the esophagus biopsy and fetal esophageal tissue. Data represents mean ±SEM (n>3 technical replicates for each biological cell line) **p<0.01, *** p<0.001 by unpaired two tailed student’s t test. **G)** Immunofluorescence staining for KRT4, KRT13 and INV showing a positive expression in all generated esophagus organoids **H)** Dual immunofluorescence staining for INV and p63 showing a basal proliferative layer positive for p63 and a suprabasal layer positive for INV, a specific marker for stratified esophagus epithelium. Scale bar 50um.

### 7. Abnormal NKX2.1 expression is retained in EA/TEF patient-derived organoids

NKX2.1 is not normally expressed in human esophagus biopsies (**Fig. 6A**). However, our patient-derived esophageal organoids showed a positive expression of NKX2.1 at levels close to the fetal trachea. As expected, no expression was observed in healthy esophageal organoids just like fetal esophagus and esophagus epithelial biopsies (**Fig. 6B**). NKX2.1 expression was interspersed in the KRT13 expressing suprabasal layers of the patient-derived esophageal organoids (**Fig. 6C, D**). A similar observation was made in TEF tissue from EA patients which showed abnormal expression of NKX2.1 (Brosens et al. 2020).

**Figure 6:**
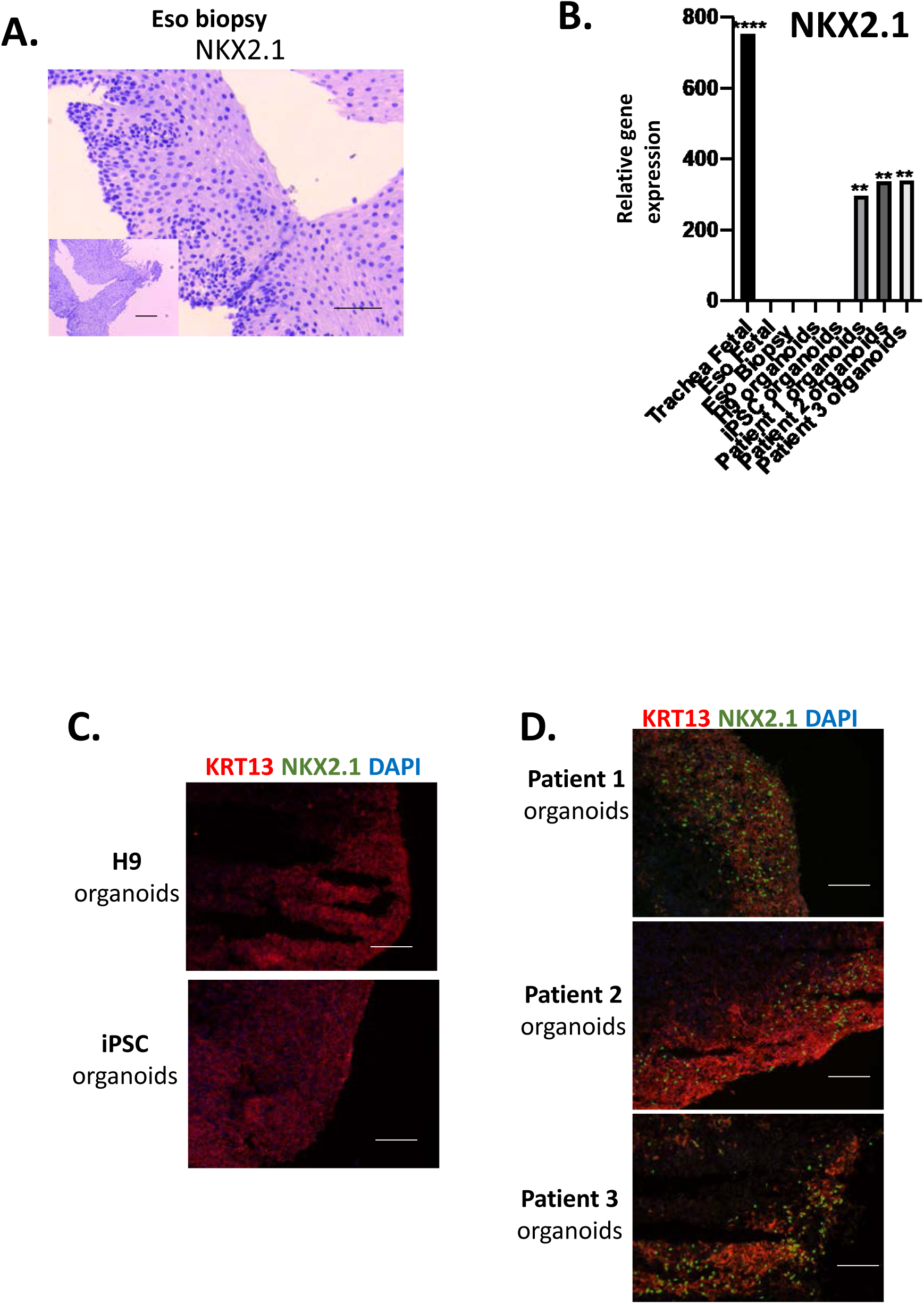
Retained abnormal expression of NKX2.1 in patient-derived esophagus organoids reveals a similarly to fistulas connecting esophagus to trachea. **A)** Immunohistochemistry of esophagus biopsy against NKX2.1 **B)** Transcript levels of NKX2.1 is significantly higher in all 3 patient-derived esophagus organoids. Expression of NKX2.1 was absent in healthy derived organoids and esophagus biopsies. Transcript levels were compared to those of H9 derived organoids. Fetal trachea was included as a reference in our analysis. Data represents mean ±SEM (n>3 technical replicates for each biological cell line) **p<0.01, *** p<0.001 by unpaired two tailed Student’s t test, **C and D)** Dual staining by Immunofluorescence confirms positive KRT13 staining in both groups, however, NKX2.1 is expressed in all 3 patient-derived esophagus organoids and is absent in healthy organoids. Scale bar 50um.

### 8. Differentiation propensity of EA/TEF iPSCs into different organ lineages is like healthy iPSCs

To verify whether the dysregulation of SOX2 at the anterior foregut stage and the abnormal expression of NKX2.1 at the mature esophageal epithelial stage in patient-derived iPSCs is specific to the esophageal fate, we investigated whether EA/TEF patients-derived iPSCs could be differentiated into other organ lineages such as tracheal, liver, and muscle progenitor cells. By using published protocol (Huang et al. 2014), we differentiated PSCs into the ventral anterior foregut cells, thereby favoring a tracheal fate, which was confirmed by the expression of NKX2.1 in both the groups **(Supplementary Fig. S3A, B).**

We then differentiated patient-derived iPSCs into posterior foregut and then directed it towards the hepatic stage. All 3 EA/TEF patient-derived iPSCs generated hepatoblast cells. Patient-derived hepatoblasts expressed Alpha fetoprotein (AFP) **(Supplementary Fig. S4).**

Finally, we directed the differentiation of the patient-derived iPSCs toward the mesodermal cell stage for generating skeletal muscle progenitor cells using a previously published protocol (Shelton et al. 2016). Myogenic progenitor cells derived from both healthy and patient iPSCs expressed similar levels of PAX3 and PAX7, which are required for myogenic specification **(Supplementary Fig. S5).** These results therefore suggest that any abnormal expression of key factors is intrinsic to the esophagus and not any other organ as is observed in these type C EA/TEF patients.

## Discussion

We report here the first *in vitro* generated matrix- and xenogeneic-free 3-dimensional mature stratified squamous esophageal epithelial organoids from EA/TEF patient-derived iPSCs. We observed a significant down regulation of SOX2 mRNA and protein expression in patient-derived anterior foregut cells. Studies have shown that SOX2 downregulation is linked to abnormal foregut separation resulting in EA/TEF (Teramoto et al. 2020; Trisno et al. 2018; Ioannides et al. 2010; Que et al. 2007; Domyan et al. 2011). We also observed an abnormal expression of NKX2.1 in patient-derived cells at the esophageal epithelial stage until the end of the organoid cultures. We also observed a distinct transcript expression profile in all 3 patient-derived anterior foregut cells, the most critical developmental time-point where patterning and subsequent separation into the esophagus and trachea occur. This dysregulation in gene and protein expression was specific to the dorsal side of the anterior foregut and therefore of the esophageal fate. In fact, directed differentiation of EA/TEF iPSCs into posterior foregut derived cells (hepatoblasts) and mesodermal cells (myoblasts) revealed a similar gene and protein expression profile to the healthy group.

The downregulation of SOX2 specifically, however, was temporary and expression levels become similar in both groups when cells were further differentiated into the mature esophageal epithelium. The exact mechanisms regulating SOX2 expression in our patient-derived cells remain unclear. Though NKX2.1 and SOX2 are hypothesized to be co-repressive master regulators of foregut separation, NKX2.1 mRNA and protein levels remained unaffected at the anterior foregut stage. Interestingly, following nanopore sequencing we observed unannotated long non-coding RNA (lncRNA-21751) lying upstream of SOX2 promoter that is significantly downregulated in all 3-patient derived anterior foregut cells. The exact role of lncRNA-21751 in regulating SOX2 expression at the anterior foregut stage remains unknown. We also speculated on the potential role of the long non-coding RNA SOX2OT at this critical stage. SOX2OT harbors the intronic region of SOX2 gene. It plays a positive role in regulating SOX2 expression in a mechanism that remains largely unknown (Shahryari et al. 2014). So, we performed qPCR analysis of SOX2OT on the anterior foregut cells and observed its expression to be downregulated in all 3 EA/TEF anterior foregut cells **(Supplementary Fig. S6).** This down regulation of the SOX2OT could be one of the regulatory molecules involved in the expression of SOX2 in the anterior foregut cells.

NKX2.1 is normally absent in the esophagus epithelium. However, in the patient esophagus epithelium and organoids, NKX2.1 mRNA and protein levels were significantly high. ISL1, a recently identified transcription factor that regulates the expression of NKX2.1, was found at similar levels in both groups **(Supplementary Fig. S1B).**

Despite relatively shallow cDNA sequencing and the presence of a batch effect overlapping two experimental variables (biological sex and sample preparation date), we identified around 173 RNA transcript isoforms that were significantly differentially expressed between the healthy and patient groups. GSTM1 was one of the most differentially downregulated genes with its distinct functions in the detoxification of electrophilic compounds including carcinogens, therapeutic drugs, environmental toxins, and products of oxidative stress. GSTM1 has a non-catalytic regulatory role in apoptotic ASK1-MAPK (mitogen-activated protein kinase) signaling cascade (Cho et al. 2001). Under non-stimulated conditions, GSTM1 inhibits apoptotic cell death (Cho et al. 2001). There has been an increasing trend of linking xenobiotics to genes involved in detoxification in early embryonic development and specifically to EA/TEF. It is suspected that an altered detoxification process triggers an alteration of proliferation or apoptotic cellular behavior that may directly affect the separation process of the foregut into the esophagus and trachea (Filonzi et al., 2010). Another interesting gene which was differentially expressed was from the family of small GTPases, Rab, which are key regulators of intracellular membrane trafficking. In a recent study, Rab11 was shown to have a direct link to epithelial remodeling and extracellular matrix degradation during the foregut separation (Nasr et al. 2019). The work shown in xenopus and mouse demonstrates how the disruption of Rab11-mediated epithelial remodeling results in tracheoesophageal clefts (Nasr et al. 2019), providing a potential mechanistic framework for foregut separation in humans. In our patient-derived anterior foregut cells, however, we observe a significant downregulation of another Rab protein, RAB37. Rab37 is a critical regulator of vesicle trafficking and play a potential role during human foregut compartmentalization, similar to what was observed in xenopus and mice with Rab11. In a suggested mechanism, Rab37 mediates exocytosis of SFRP1 (secreted frizzled related protein 1) an antagonist of the Wnt pathway to suppress Wnt signaling in lung cancer cells *in vitro* (Cho et al. 2018). Knowing the particular importance of the inhibition of Wnt signaling in the anterior foregut to favor an esophageal fate (Woo et al. 2011), raises the potential role of Rab37 at this developmental stage. Furthermore, the identification of numerous new transcript isoforms, including known and novel long non-coding RNAs, supports the observed regulatory complexity of esophagus and trachea development, as well as EA/TEF etiology, whilst suggesting that non-coding regulatory transcripts play a role in this process.

We cannot exclude a role of mesenchymal cells in the dysregulation of SOX2 and NKX2.1 in the present experimental setting. We detected mRNA expression of brachyury, a transcription factor that regulates mesoderm formation (Herrmann et al. 1990) and vimentin, an intermediate filament expressed in mesenchymal cells, in our cultures during directed esophagus differentiation **(Supplementary Fig. S7A).** Additionally, after 2 months of culture, we observed vimentin protein expression in our mature esophageal organoids derived from both healthy and patient cells **(Supplementary Fig. S7B).** The influence of mesenchyme on foregut epithelium division has been previously demonstrated (Han et al. 2020). Dysregulation of SOX2 has been linked to mesenchymal development with respiratory characteristics (Teramoto et al. 2020).

In conclusion, the experimental approach of using EA/TEF derived iPSCs, allowed us to mimic the initial developmental stages of the human esophagus to understand the origins of this malformation. We can conclude that the intrinsic defect observed in these cells are limited only to the esophagus. Our work is limited to isolated type C EA/TEF and thus cannot relate our results to other types of EA or syndromic EA with associated malformations. More studies are required to understand the failure of key mechanisms and pathways involved during the critical interface of anterior foregut specification. This work therefore highlights the importance of using patient-derived iPSCs to model congenital diseases to yield new insights on organ development during embryogenesis.

## Materials and methods

### Blood collection for reprogramming

Blood was collected from the three pediatric patients after obtaining consent from their parents to reprogram the blood cells to PSCs for research purposes. This study was approved by the Institutional Review Board of CHU-Sainte Justine Research Center (Protocol #2018-1670 (For iPSCs); 2019-2102 (For WES).

### Experimental Design

Using the Institutional iPSC Core facility, we reprogrammed PBMCs from three different EA/TEF type C patients. Control and patient cells were not matched for sex and ethnicity.

All iPSCs used for differentiation was between passages 20 and 35. Healthy and patient-derived iPSCs were differentiated simultaneously into mature esophageal organoids for every directed esophageal differentiation. The identity of the samples was not blinded to the investigator. Hepatic and myoblast differentiation were performed in Paganelli and Dumont labs, respectively. iPSCs derived from healthy subjects for esophageal differentiation, hepatic differentiation, and myoblast differentiation are different. The same clones of EA/TEF patient-derived iPSCs were used for all directed differentiations.

### Human embryonic stem cell and induced pluripotent stem cells

Human embryonic stem cell (ESC) cell line, H9 was a kind gift from the Andelfinger lab at CHU Sainte Justine Research Center (Wunnemann et al. 2020). Healthy iPSC cell line (GHC4) and EA/TEF patient-derived iPSCs (EA1, EA2, and EA3) were generated and obtained from the IPSC Core Facility at CHU-Sainte Justine research center.

### Culture and expansion of ESCs and iPSCs

Both ESCs and iPSCs were cultured on feeder free and non-xenogeneic conditions. Cells are plated on human vitronectin VTN XF^TM^ (STEMCELL Technologies, Canada. Catalog#100-0763) coated 100mm cell culture dishes. Cells were maintained at 37°C with 5% CO_2_ with daily replacement of Essential 8 media system (Thermofisher, Canada. Catalog#A1517001). Cells were passaged as aggregates every 3-4 days until they reach 60-70% confluency with 0.5mM EDTA diluted in PBS (Thermofisher, Canada. Catalog#15575020).

### Differentiation protocol-Preparing cells for differentiation (Day -1 and Day 0)

Two days prior to differentiation (D-1), cells were dissociated into single cells using Accutase^TM^ (Stem Cell technologies, Canada. Catalog# 07922) and transferred onto BIOLAMININ 521 LN (LN521, BioLamina, Sweden. Product# LN521-02) coated plates with E8 media supplemented with 10uM Rock inhibitor Y-27632 (Sigma-Aldrich, US. Product# SCM075). The following day (D0) (12-16 hours later), media was changed with E8 only. If survival rate of the cells were less than 50%, they were cultured for an additional 24 hours before starting the differentiation. ESC and iPSCs were maintained at 37°C with 5% CO_2_ throughout the differentiation process.

### Endoderm differentiation (Day 1 to Day 3)

We modified and adapted previously published protocol for endoderm differentiation (Matsuno et al. 2016). Xeno-free media (XFM-) was prepared using 500 mL of RPMI 1640 medium without L-Glutamine (Thermo Fisher Scientific, Canada. Catalog# 11875101), 10mL B-27 Supplement minus insulin (Thermo Fisher Scientific, Canada. Catalog# A1895601), 5mL GlutaMax^TM^ (Thermo Fisher Scientific, Canada. Catlog# 35050061), 5mL of KnockOut^TM^ serum replacement (Thermo Fisher Scientific, Canada. Catalog# 10828010), 5mL Penicillin-Streptomycin (10,000 U/mL) (Thermo Fisher Scientific, Canada. Catalog# 15140148), 7.5mL HEPES (1M) Buffer (Thermo Fisher Scientific, Canada. Catalog# 15630130), 5mL of MEM Non-essential amino acids (100X) ((Thermo Fisher Scientific, Canada. Catalog# 11140050). Day 1 of differentiation cells are first washed with XFM-media to remove any residual E8 media, then cultured in XFM- with 100ng/mL Activin A (R&D systems, US. Catalog# 338-AC-010/CF) and 3uM CHIR99021 (Stem Cell technologies, Canada. Catlog# 72052). On Days 2 and 3 of culture, cells were first washed with XFM-media and the culture was continued with XFM-supplemented with 100ng/mL Activin A and 250nM of LDN193189 (Stemgent, US. Code# 04-0074). By day 3, a 70-80% confluent monolayer of endodermal cells should be observed under the microscope.

### Anterior foregut differentiation (Day 4 and Day 5)

Endodermal cells were first washed with XFM- media and then cultured for 24 hours in XFM- supplemented with 1uM A8301 (Stemgent, US. Code# 04-0014) and 250nM of LDN193189. The following day cells were washed XFM- media and cultured in XFM- supplemented with 1uM A8301 and 1uM IWP2 (Stemgent, US. Code# 04-0034).

### Esophagus differentiation (Day 6 to Day 24)

From day 6 to day 16 we switch to XFM+ containing the same components as XFM- and replacing b27 supplement minus insulin with b27 supplement with insulin (Thermofisher, Canada. Catalog# 17504044). To induce esophageal fate, we modified and adapted previously published protocol (Zhang et al. 2018; Trisno et al. 2018). Anterior foregut cells were cultured in XFM+ supplemented with 1uM A8301 and 250nM LDN193189 from day 6 until day 16, changing media daily. By day 16, we observed that cells had reached 100% confluency and the presence of dense cell clusters. On day 16 esophageal progenitor cells are cultured in XFM+ only until day 24.

### Esophageal Organoid formation (2 months)

Organoids were generated in suspension using Nunclon^TM^ Sphera^TM^ low attachment 96-well plates (Thermofisher, Canada. Catalog# 174930) by modifying previously published esophageal studies (Giroux et al. 2017; DeWard, Cramer, and Lagasse 2014). On day 24 of esophageal differentiation, cells were detached using TrypleE (Thermo Fisher Scientific, Canada. Catalog# 12604013) and gently resuspended in XFM+ media. Viable cells were counted using Trypan blue solution (Thermofisher, Canada. Catalog# 15250061) and 50,000 cells are then aliquoted in each well of the 96-well plate containing XFM+ supplemented with 1uM A8301, 250nM LDN, 3uM CHIR99021, 20ng/mL FGF2/bFGF (Peprotech, US. Catalog# AF100-18B) and 200ng/mL EGF (Thermofisher, Canada. Catalog# PHG0313).

### Other organ-lineage differentiation

#### - Trachea differentiation (Day 6 to Day 16)

To induce tracheal fate, we again modified previously published protocols (Huang et al. 2014; Huang et al. 2015). Anterior foregut cells were cultured from day 6 to day 16 in XFM+ supplemented with 3uM CHIR99021, 10ng/mL human FGF10 (R&D systems, US. Catalog# 345-FG-025/CF), 10ng/mL human FGF7 (R&D systems, US. Catalog# 251-KG-010/CF), 10ng/mL BMP4 (Peprotech, US. Catalog# 120-05) and 50 nM RA (Tocris, UK. Catalog# 0695) from day 6 to day 16 changing media daily. Cells reached 100% confluency and formed two-layered cell clusters.

#### - Hepatoblast differentiation (Day 0 to day 15)

iPSCs were dissociated by TrypleE (Life Technologies, Catalog# 12604013) to single cells and seeded on human recombinant laminin 521 (Biolamina)-coated plates in Essential 8 Flex medium at a density of 7×10^^5^ cells/cm^2^. Differentiation starts (day 0) when the cells reach around 70% confluency by changing the medium to RPMI B27 minus insulin (Life technologies) supplemented with 1% knockout serum replacement (KOSR, Life technologies, Catalog# 10828010). For the first 2 days, the cells were exposed to 100ng/mL Activin A (R&D systems, Catalog# 338-AC-010/CF) and 3uM CHIR99021 (Stem Cell Technologies, Catalog #72052) and then for the following 3 days to 100ng/mL Activin A alone. Subsequently, RPMI B27 minus insulin medium was supplemented with 20ng/mL BMP4 (Peprotech, Catalog# 120-05), 5ng/mL bFGF (PeproTech, Catalog# AF100-18B) and 4uM IWP2 (Tocris, Catalog# 3533) and 1uM A83-01(Tocris, Catalog# 2939/10) for 5 days, with daily medium change. At day 10, the medium was changed to RPMI B27 (Life technologies, Catalog# 17504044), supplemented with 2% KOSR, 20ng/mL BMP4, 5ng/mL bFGF, 20ng/mL HGF (PeproTech, Catalog #100-39H) and 3uM CHIR99021 for 5 days, with daily medium change.

#### - Myoblast differentiation (Day 0 to Day 20)

The generation of iPSC-derived myoblast was adapted from a protocol published by M. Shelton *et al* with minor modifications (Shelton et al. 2016). Briefly, different growth factors and inhibitors are sequentially used to drive iPSCs toward the mesodermal lineage and promote their myogenic cell fate commitment. One day prior differentiation, human iPSCs (patients and control) were dissociated with TrypLE™ (Gibco, Catalog# 12604013) and 10^5^ cell/well were plated as small colonies (10-20 cell/colony) on Vitronectin-coated 12-well plates using mTeSR1 media (STEMCELL Technologies, Catalog# 12604013) supplemented with 10 μM ROCK inhibitor (Y-27632, STEMCELL Technologies, Catalog# 12604013). Next day, the medium was changed to TeSR-E6 media (STEMCELL Technologies, Catalog#05946) supplemented with 7 μM CHIR 99021(STEMCELL Technologies, Catalog #72052) for 3 days. After three days of CHIR99021treatment, cells were gently washed with DPBS and cultured only in TeSR-E6 medium without any CHIR99021, and media was changed every day till day 7. At this time point, a borad expression of the somite markers PAX3 and MEOX1 can be detected. From days 10 to 20 of differentiation, 5ng/mL FGF2 (Wisent) is added to the TeSR-E6 medium to promote myogenic cell proliferation. At day 20, a significant proportion of cells express the muscle stem cell marker PAX7.

### RT-qPCR

At each developmental stage (definitive endoderm, anterior foregut, esophageal progenitors, mature esophageal epithelium/organoids and other organ-lineages), cells were detached using Accutase^TM^ and RNA was extracted using the Promega kit ReliaPrep ^TM^ RNA Cell Miniprep System (Promega, US. Catalog# Z6011). RNA was reverse transcribed using Omniscipt ^TM^ RT kit (Qiagen, US. Catalog# 205113) and the complementary DNA (cDNA) obtained was used for real time quantitative PCR using LightCycler instrument (Roche Life Science, Germany). cDNA was quantified using TaqMan Gene expression assays and the TaqMan primers to target genes were purchased from Thermo Fisher Scientific listed in **Table 1**. The transcript level of each gene was normalized to GAPDH housekeeping gene using the 2^-DDCT method. Relative gene expression was calculated and reported as fold change compared to indicated samples using GAPDH normalized transcript level. The results include a mean of at least 3 technical replicates for each biological sample.

**Table 1:** List of Taqman gene expression assays for qPCR analysis.

| Gene name | Source | Identifier |
| --- | --- | --- |
| CXCR4 | ThermoFisher | Hs00607978_s1 |
| GATA4 | ThermoFisher | Hs00171403_m1 |
| SOX17 | ThermoFisher | Hs00751752_s1 |
| SOX2 | ThermoFisher | Hs04234836_s1 |
| PAX9 | ThermoFisher | Hs00196354_m1 |
| ISL1 | ThermoFisher | Hs00158126_m1 |
| NKX2.1 | ThermoFisher | Hs00968940_m1 |
| KRT4 | ThermoFisher | Hs00361611_m1 |
| KRT13 | ThermoFisher | Hs02558881_s1 |
| P63 | ThermoFisher | Hs00978340_m1 |
| INV | ThermoFisher | Hs00846307_s1 |

**Table 2:** List of primary and secondary antibodies.

| Antibodies | Source | Identifier |
| --- | --- | --- |
| Mouse monoclonal antibody to TFF1 | Abcam | ab72876 |
| Rabbit monoclonal antibody to CXCR4 | Abcam | ab181020 |
| Mouse monoclonal antibody to SOX2 | Abcam | ab79351 |
| Rat monoclonal antibody to PAX9 | Abcam | ab28538 |
| Rabbit monoclonal antibody to cytokeratin 4 | Abcam | ab51599 |
| Mouse monoclonal antibody to SOX17 | Abcam | ab84990 |
| Rabbit monoclonal antibody to ISL1 | Abcam | ab109517 |
| Rabbit polyclonal antibody to GATA4 | ThermoFisher | PA1-102 |
| Mouse monoclonal antibody to INV | Abcam | ab68 |
| Rabbit polyclonal antibody to P63 | Abcam | ab53039 |
| Rabbit monoclonal antibody to KRT13 | Abcam | ab92551 |
| Rabbit polyclonal antibody to AFP (ready to use) | DAKO omnis | GA50061-2 |
| **Secondary antibodies** |  |  |
| Goat anti-Mouse IgG H&L (Alexa Fluor 488) | Abcam | ab150113 |
| Goat anti-Mouse IgG H&L (Alexa Fluor 594) | Abcam | ab150120 |
| Donkey anti-Rabbit IgG H&L (Alexa Fluor 488) | ThermoFisher | A21206 |
| Goat anti-Rabbit IgG H&L (Alexa Fluor 488) | Abcam | ab150077 |

### Immunofluorescence and microscopy imaging

For immunofluorescence staining, cells at each developmental stage (definitive endoderm, anterior foregut, esophageal progenitors, mature esophageal epithelium/organoids, and other organ-lineages) were fixed with 4% paraformaldehyde (PFA) (Thermo Scientific, Canada. Catalog# AAJ19943K2) for 20 minutes at room temperature then washed 3 times with DPBS. Cells were then permeabilized with 0.4% Triton X-100^TM^ (Sigma-Aldrich, US. Catalog# 9002-93-1) in DPBS for 25 minutes at room temperature followed by washing with DPBS. Cells were then incubated with 3% blocking serum in DPBS for 1 hour at room temperature. Primary antibody was added to antibody dilution buffer (1% PBS-BSA, 0.3% Triton X-100, 0.3% serum) and were incubated overnight at 4°C. The following day, the cells were washed 3 times with DPBS. Secondary antibody was diluted in the same antibody dilution buffer, added to cells and incubated for 1 hour at room temperature (primary and secondary antibodies listed in **Table 3**). Following washing with DPBS, cells were stained with DAPI for 15 minutes at room temperature. Cover slips are mounted on top of a drop (7-8 uL) of ProLong^TM^ Diamond Antifade Mountant (Thermo Fisher Scientific, Canada. Catalog# P36970).

### RNA-sequencing assay

**RNA was extracted as described in the RT-qPCR section which was followed by** library preparation for Nanopore Sequencing was done using two different protocols. RNA (50 ng total RNA) was spiked reverse transcribed and amplified by PCR following the manufacturer’s instructions for the cDNA-PCR Sequencing kit (SQK-PCS109, [Oxford Nanopore Technologies, UK) up until the PCR step (14 cycles, 500 seconds extension). The PCR reactions were then prepared for sequencing using the Genomic DNA by Ligation kit (SQK-LSK109 [Oxford Nanopore Technologies, UK]). Concentrations were quantified for RNA after elution and for cDNA after the PCR step and before loading the flow cells using a Qubit Fluorometer with a Qubit RNA assay kit (High sensitivity) for RNA and a Qubit dsDNA assay kit (Broad range) for cDNA [Invitrogen, Waltham, Massachusetts, USA]. Final libraries, with the following cDNA concentrations: 26.2 ng/uL (Patient 1), 29 ng/uL (Patient 2), 23.8 ng/uL (Patient 3), 28ng/uL (iPSC), 13.4 ng/uL (H9), were loaded onto MinION flow cells (R9.4.1 - FLO-MIN006) and ran for 74 hours on GridION and MinION Mk1C sequencers. When required, the sequencing runs were refueled with 250 uL of FB buffer.

### RNA-sequencing analysis

Raw fast5 files were basecalled during the sequencing run using Guppy v4.0.11 [https://nanoporetech.com/] in high accuracy mode. Fastq_pass and fastq_fail files for each sample were submitted to Pychopper v2.4.0 [https://github.com/nanoporetech/pychopper] to identify full-length reads, split ligation concatemers, rescue fused reads and reorient reads based on the stranded barcode adapters. Full length and rescued reads were then aligned to the human reference genome (GRCh38.p13)(Frankish et al. 2019), using Minimap2 v2.18(Li 2018) with -a -x splice --MD -- secondary=no options. Alignments were converted to bam format and sorted with samtools v1.12 (Li et al. 2009). Resulting bam files were merged using “samtools merge” before being used as input for de novo assembly with Stringtie2 v2.1.4 (Kovaka et al. 2019). Gencode reference transcriptome v37 gtf format was used as input for Stringtie2’s -G option and all transcripts were collapsed with the long reads -L parameter. GFFcompare v0.1.12.2(Pertea and Pertea 2020) was used to map the resulting GTF file to Gencode to evaluate the assembly and filter it. Transcript per million (TPM), classcode and exon number filters have been applied manually to select all isoforms that are TPM>0.2, “=” or “c” classcode (provided by GFFcompare) and all classcode if more than 1 exon. Fasta sequence corresponding to the filtered assembly annotation were retrieved using GFFread v0.12.7 (Pertea and Pertea 2020) and the samples fastq sequences obtained after pychopper were aligned again using minimap2 with -k 14 -a -N 100 options and the fasta sequence from the filtered assembly as a reference. Bam files were obtained using samtools and isoforms were quantified using Salmon v1.5.2 (Patro et al. 2017) in quant mode with -l SF –noErrorModel –noLengthCorrection options as recommended for Nanopore long reads. Read counts from all samples were merged into the same matrix using Salmon quantmerge option. Isoforms with more than 3 samples with null read counts were filtered out manually to avoid artefacts. Normalization and differential expression analysis was performed using DESeq2 [Bioconductor version 3.13] (Love, Huber, and Anders 2014) on R v4.1.0 (R Core Team, 2021) to generate a normalized count matrix and statistics on differential expression. Batch effect correction was done using the sva package (Leek et al. 2021). Isoforms with a p-value < 0.01 and a |log2(FoldChange)|>1 after batch correction were considered statistically significant in the differential expression analysis. Isoforms with a p-value<0.01 and |log2 (FoldChange)|>0.5 were used to perform a GO enrichment analysis with GOrilla from Gene Ontology (Eden et al. 2009).

### Quantification and statistical analysis

All data quantification is presented as the mean ± SEM using GraphPad Software Prism 6. Statistical significance was determined by Student’s t tests, Mann-Whitney test. When more than two groups are compared, multiple comparisons were performed using one-way ANOVA to compare the two groups. For each analysis, at least 3 technical replicates of each biological cell lines were included. Representative pictures shown are indicated in the legends. P values of 0.05 or less were considered statistically significant.

## Acknowledgments

We greatly thank Dr. Basma Benabdallah, iPSC Core at CHU Sainte-Justine Research Centre and Dr. Silvia Selleri (Paganelli Lab) from CHU Sainte-Justine Research Center.

This work was funded by the Foundation CHU Sainte-Justine and the Association Quebecoise de l’atresie de L’oesophage (AQAO) to Dr. Christophe Faure.

Z.O. is supported by an award from MITACS. N.A.D. is supported by FRQS (Fonds de recherche du Québec – Santé) Junior-2 award and by research grants from the Canadian Institutes of Health Research (PJT-156408, PJT-174993), Natural Sciences and Engineering Research Council of Canada (RGPIN-2018-05979), Canada Foundation for Innovation (37622), and the Quebec Cell, Tissue and Gene Therapy Network –ThéCell (a thematic network supported by the FRQS).

MAS is supported by a FRQS Junior 1 fellowship and establishment award (295760).

## Author Contributions

AD, CF, and SR designed the study. SR performed the experiments. SR and AD went through the data analysis. MS and MS performed the nanopore RNA sequencing, data acquisition and analysis. ND and ZO were responsible for myoblast differentiation. SR, AD and CF wrote the manuscript. AD and CF reviewed the manuscript.

## Supplementary Figures

**Supplementary figure S1:**
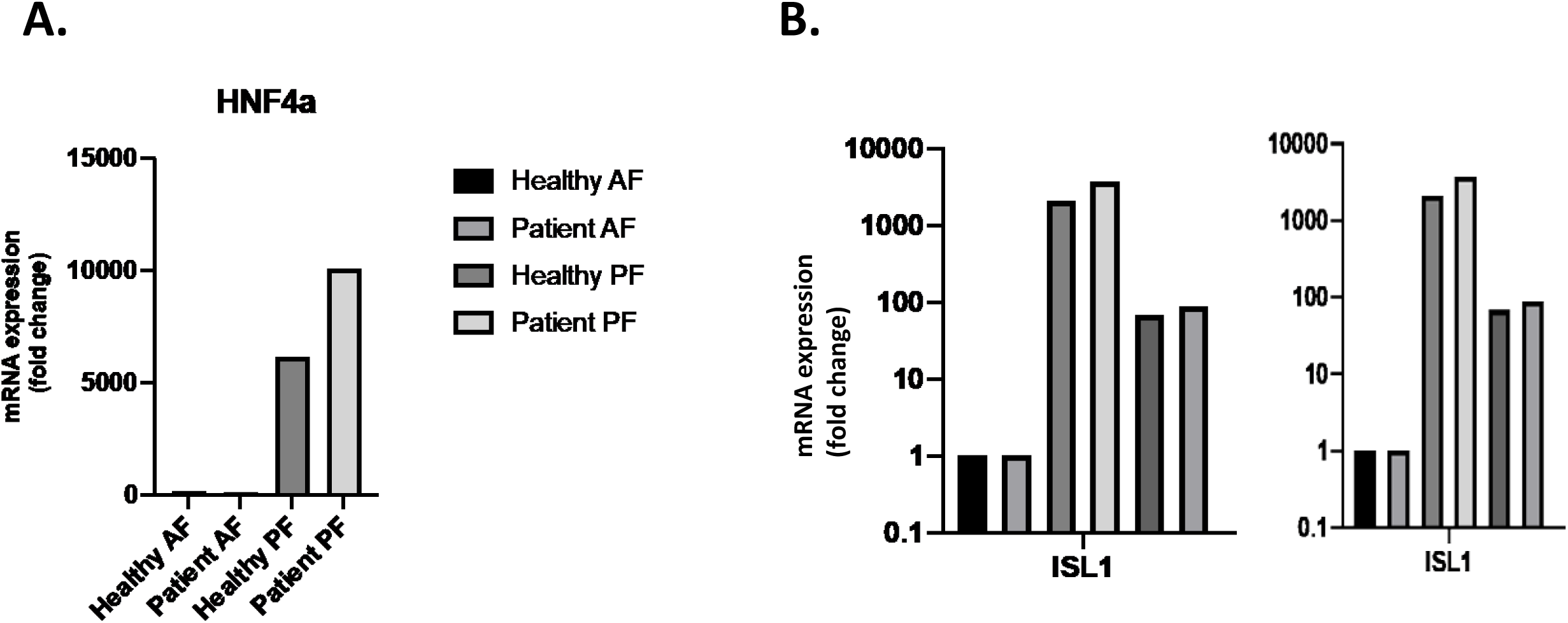
mRNA expression levels of HNF4a, ISL1 in healthy and patient derived cells. **A)** Expression levels of HNF4a, a posterior foregut marker, by qPCR is absent in healthy and patient derived anterior foregut cells compared to posterior foregut cells. mRNA levels were represented by the fold change compared to healthy posterior foregut cells**. B)** Expression of ISL1 mRNA levels by qPCR at definitive endoderm, anterior foregut and esophagus epithelium at day 24 are similar between healthy and patient cells. Fold change relative expression was compared to healthy definitive endodermal cells. **n=3 biological replicates**

**Supplementary figure S2:**
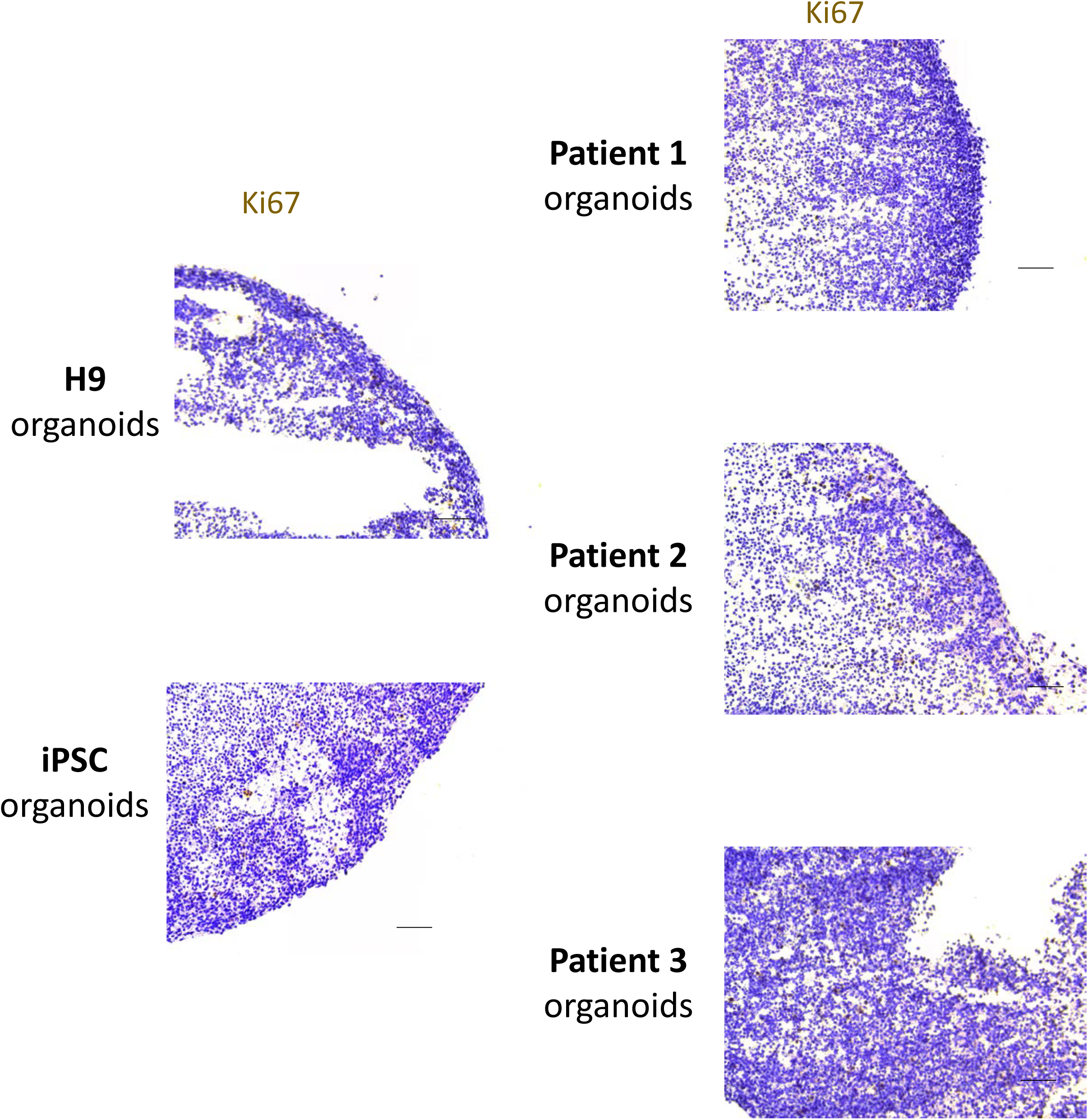
Healthy and Patient-derived organoids can proliferate after 2 months of culture. Immunohistochemical staining for ki67, a proliferative marker, shows a similar proliferative capacity in both healthy and patient derived organoids after 2 months of culture. Scale bar 50um

**Supplementary figure S3:**
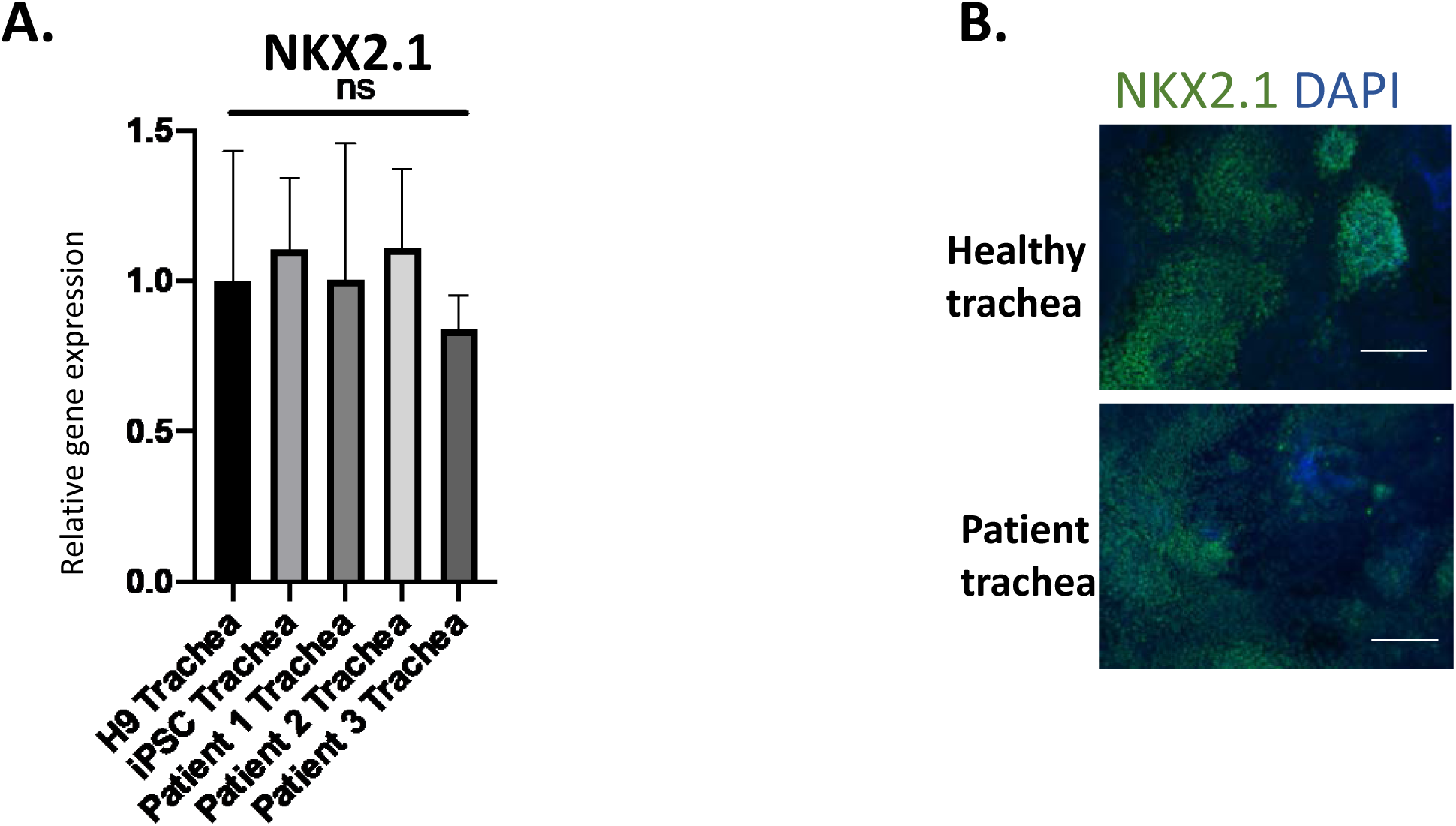
Healthy and Patient-derived iPSCs can generate tracheal epithelium. **A)** Expression of NKX2.1 transcription factor in tracheal epithelial cells derived from healthy and patient iPSCs reveal similar expression levels. mRNA relative expression was compared to the healthy group. Data represent mean +- SEM (n=3 technical replicates for each biological cell line). **B)** Immunofluorescence staining of tracheal epithelium from healthy and patient cells reveal expression of NKX2.1 in both groups. Negative controls were included for each staining. Scale bar 50um.

**Supplementary figure S4:**
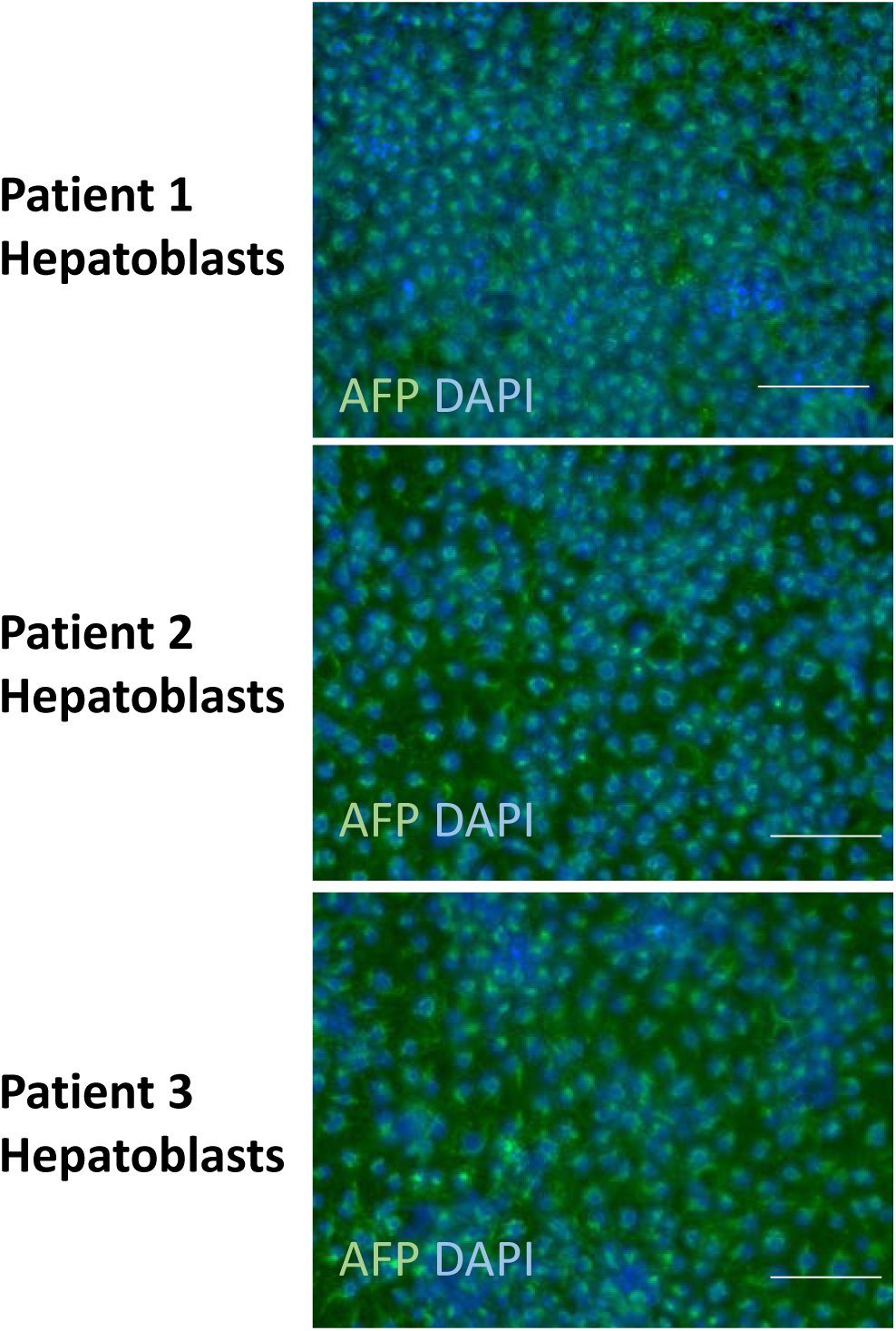
Hepatic differentiation of healthy and patient derived iPSCs. Immunofluorescence staining of all 3-patient derived hepatoblasts reveal expression of AFP a specific hepatic marker. Scale bar 50um

**Supplementary figure S5:**
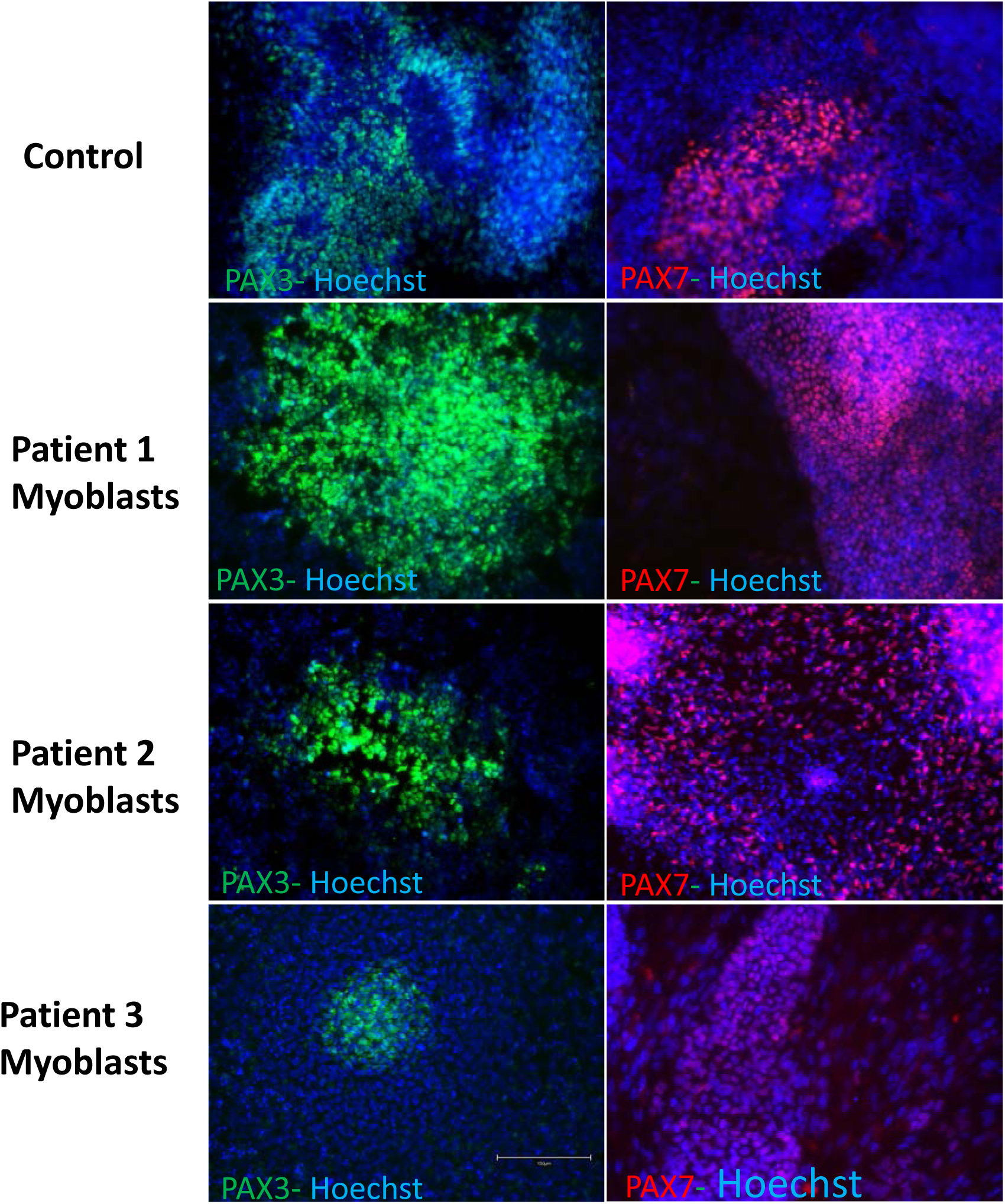
Differentiation of iPSCs into myoblast cells. Immunofluorescence staining of 3 patient-iPSC derived myoblasts reveal similar protein expression of PAX3 and PAX7 similarly to the control group. Scale bar 100um

**Supplementary figure S6:**
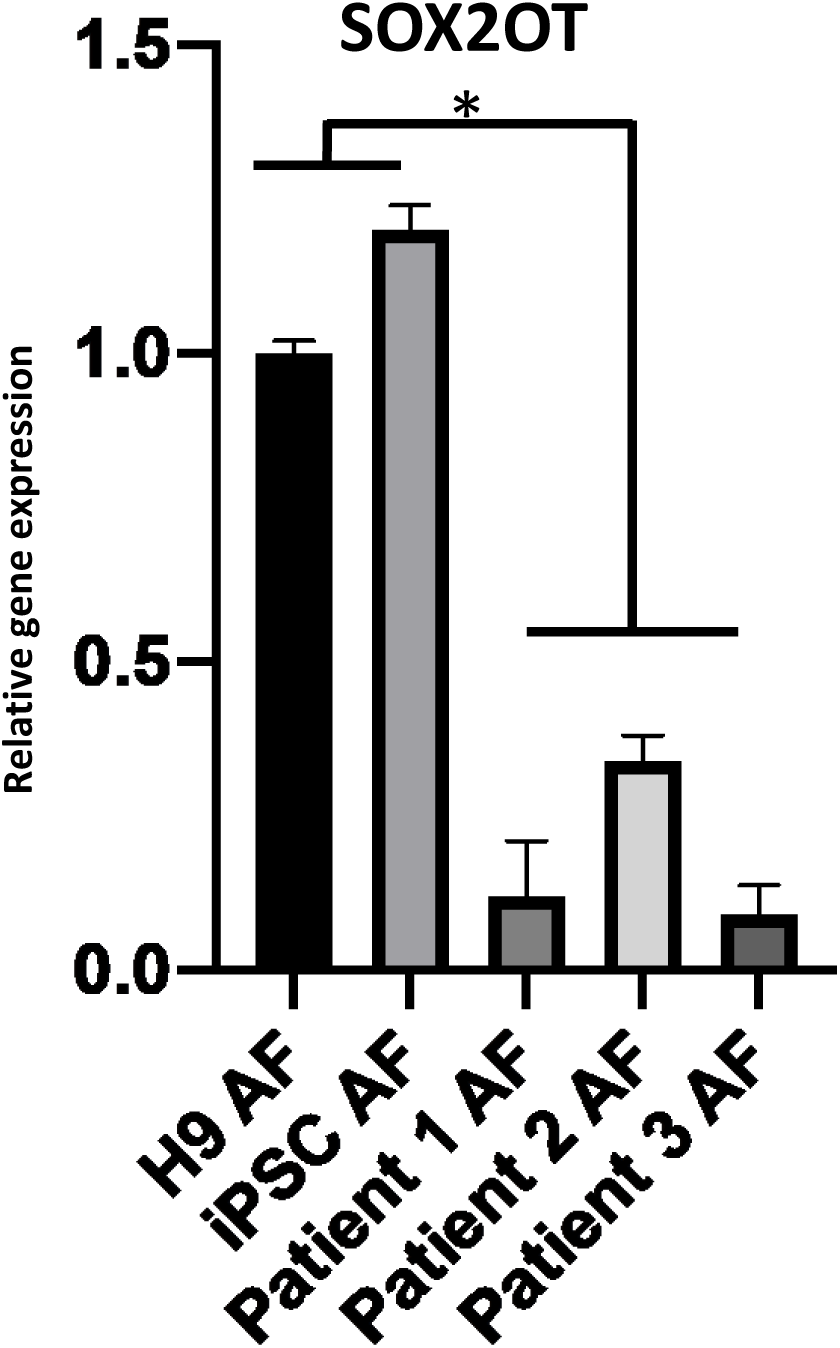
Long non-coding RNA SOX2OT a potential regulator of SOX2 at the anterior foregut stage. mRNA levels by qPCR reveal significant downregulation of SOX2OT in all 3 patient-derived anterior foregut cells. Relative expression was compared to H9 AF cells. N=3 biological replicates.

**Supplementary figure S7:**
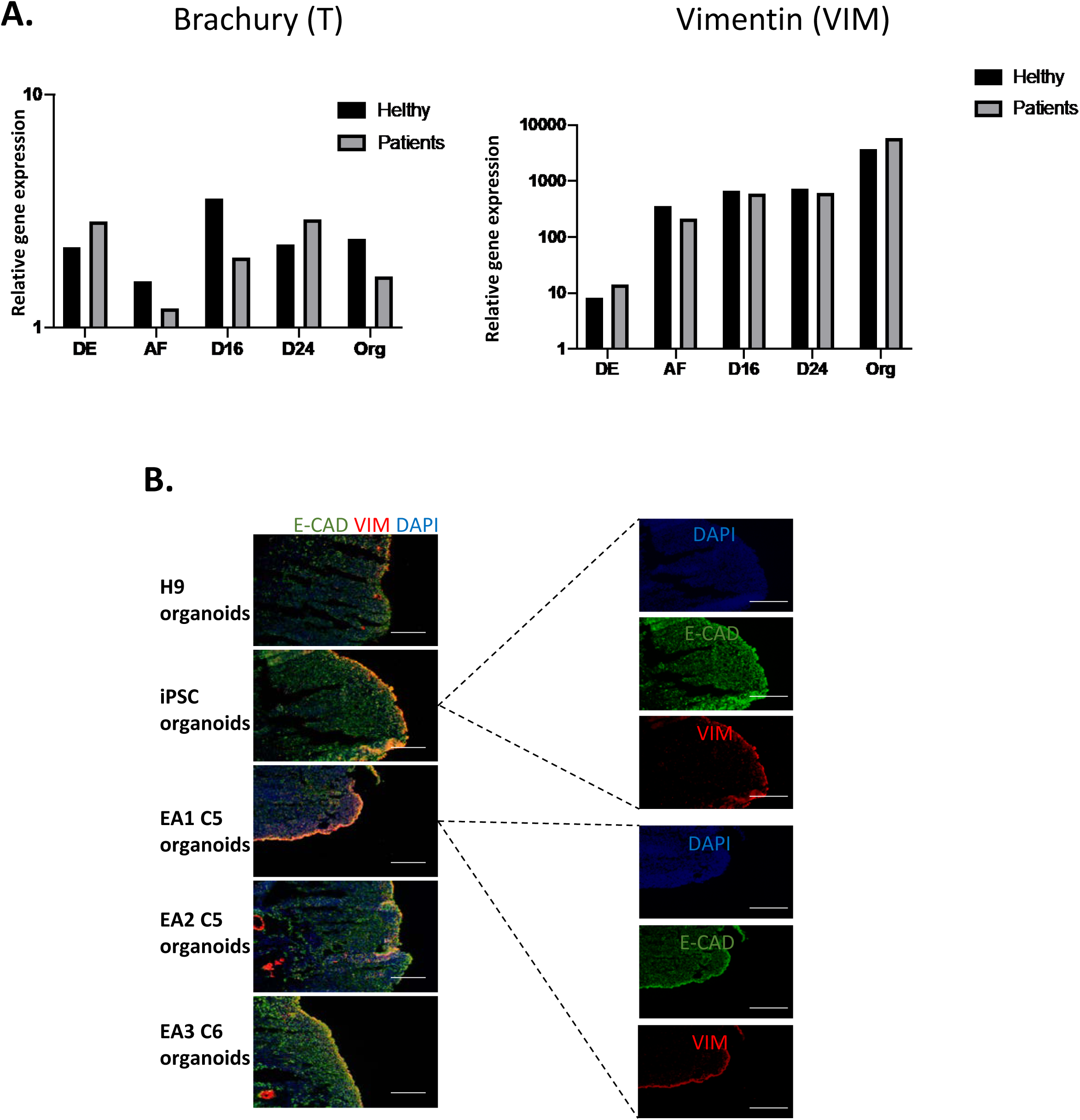
2 months mature esophagus organoids express mesenchymal markers brachyury and vimentin. **A)** Transcript levels of Brachyury (T) and Vimentin (Wim) throughout esophagus differentiation is similar in both healthy and patient derived cells. Transcript levels were compared to those of day 0 iPSCs n=3 biological replicates. B) Healthy and patient derived esophagus organoids express by immunofluorescence vimentin VIM in the outermost layer of the esophagus whereas the rest of the esophagus is positive for E-cadherin, an epithelial marker. Esophagus organoids are organized with an inner core of epithelial cells and outer layer of mesenchymal cells. Scale bar 50um

---

**Supplementary Table 1a:**
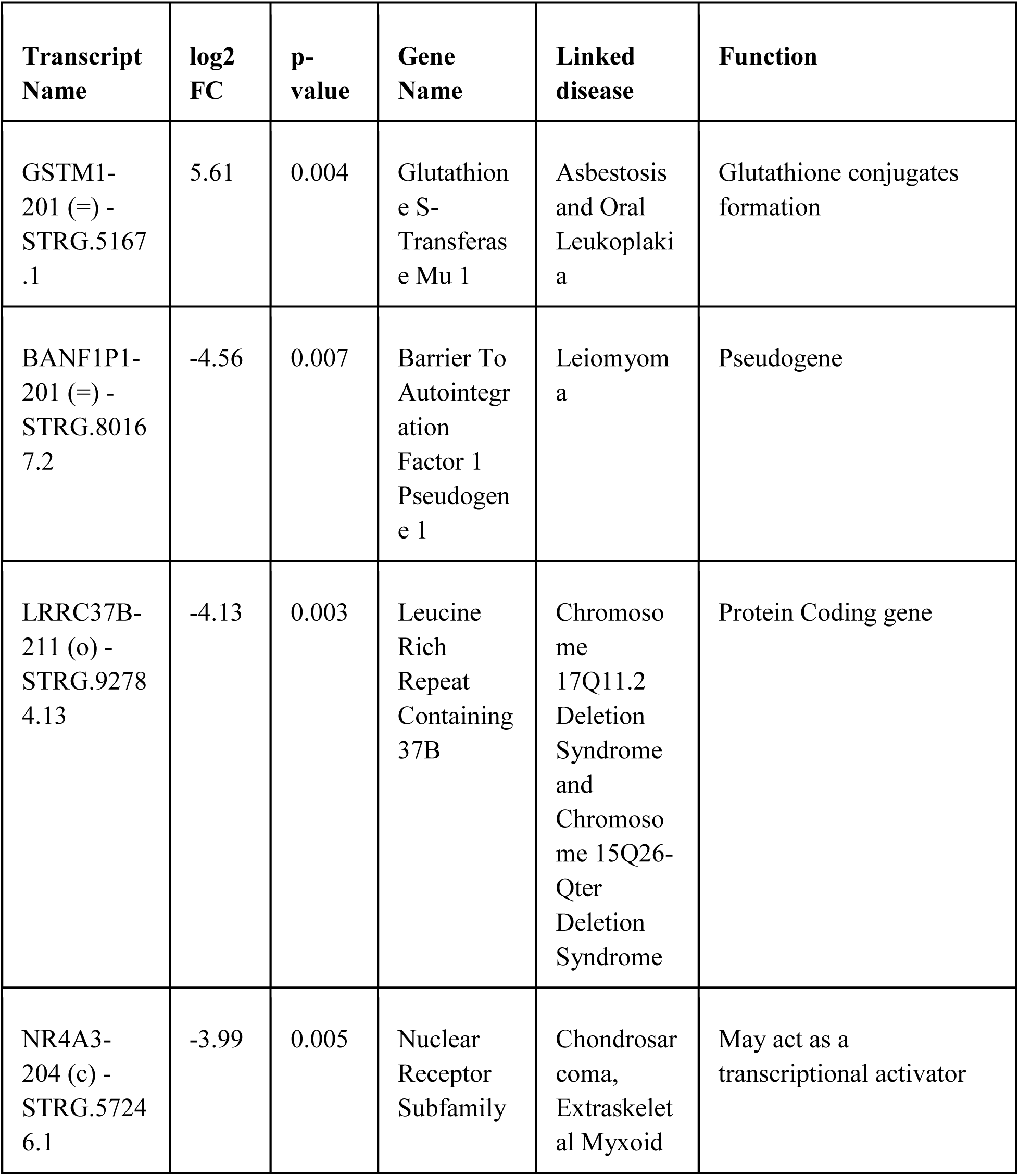
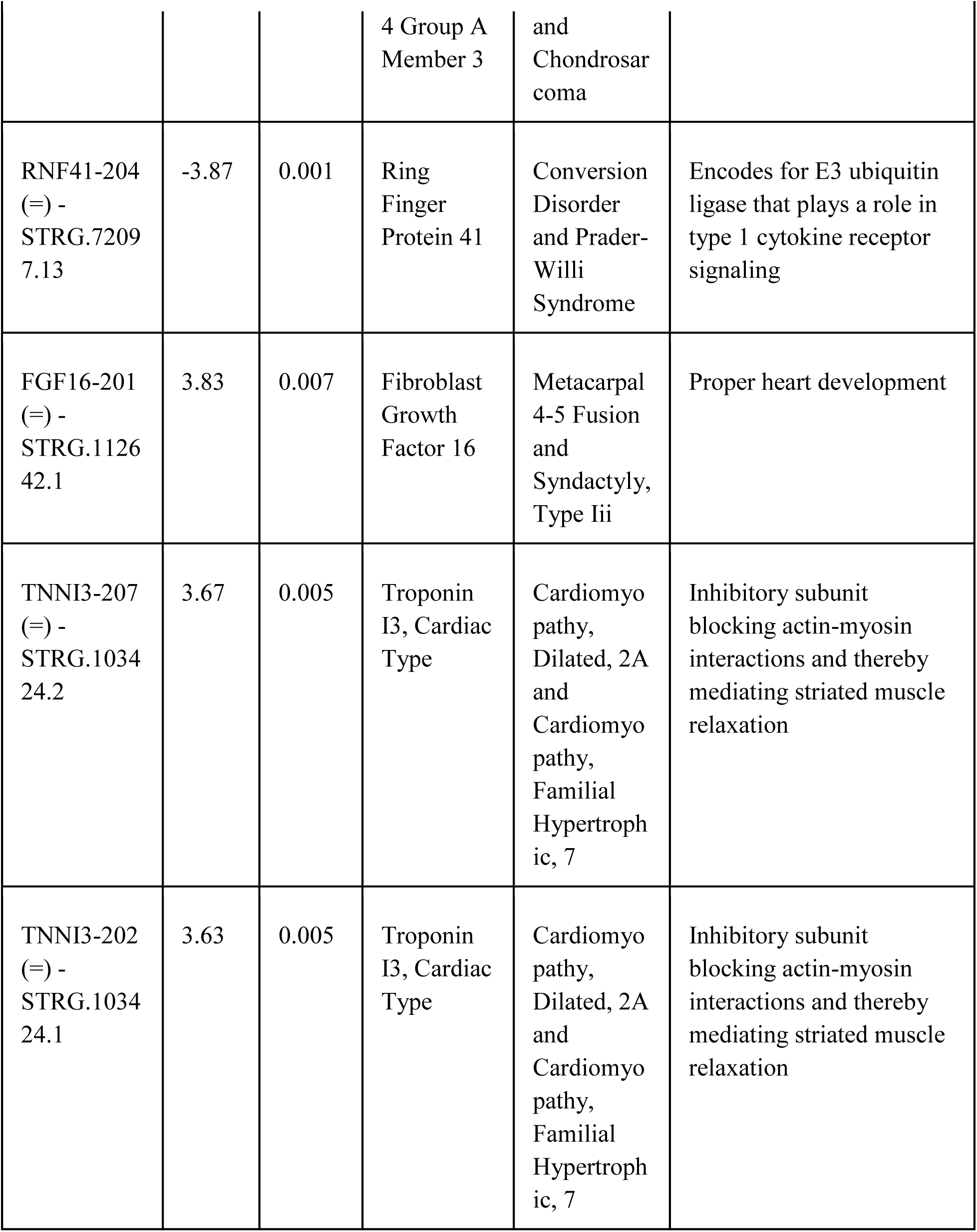
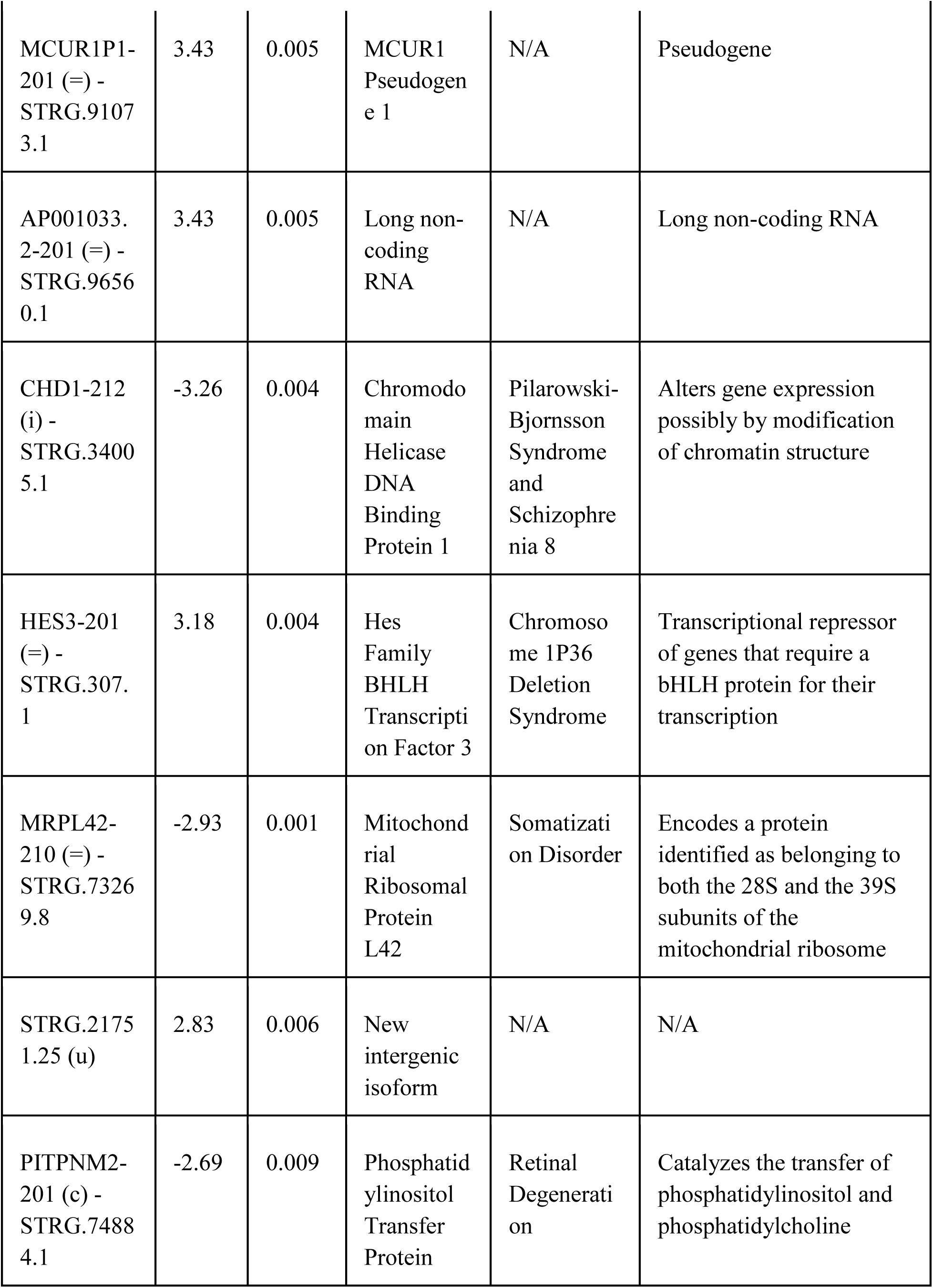
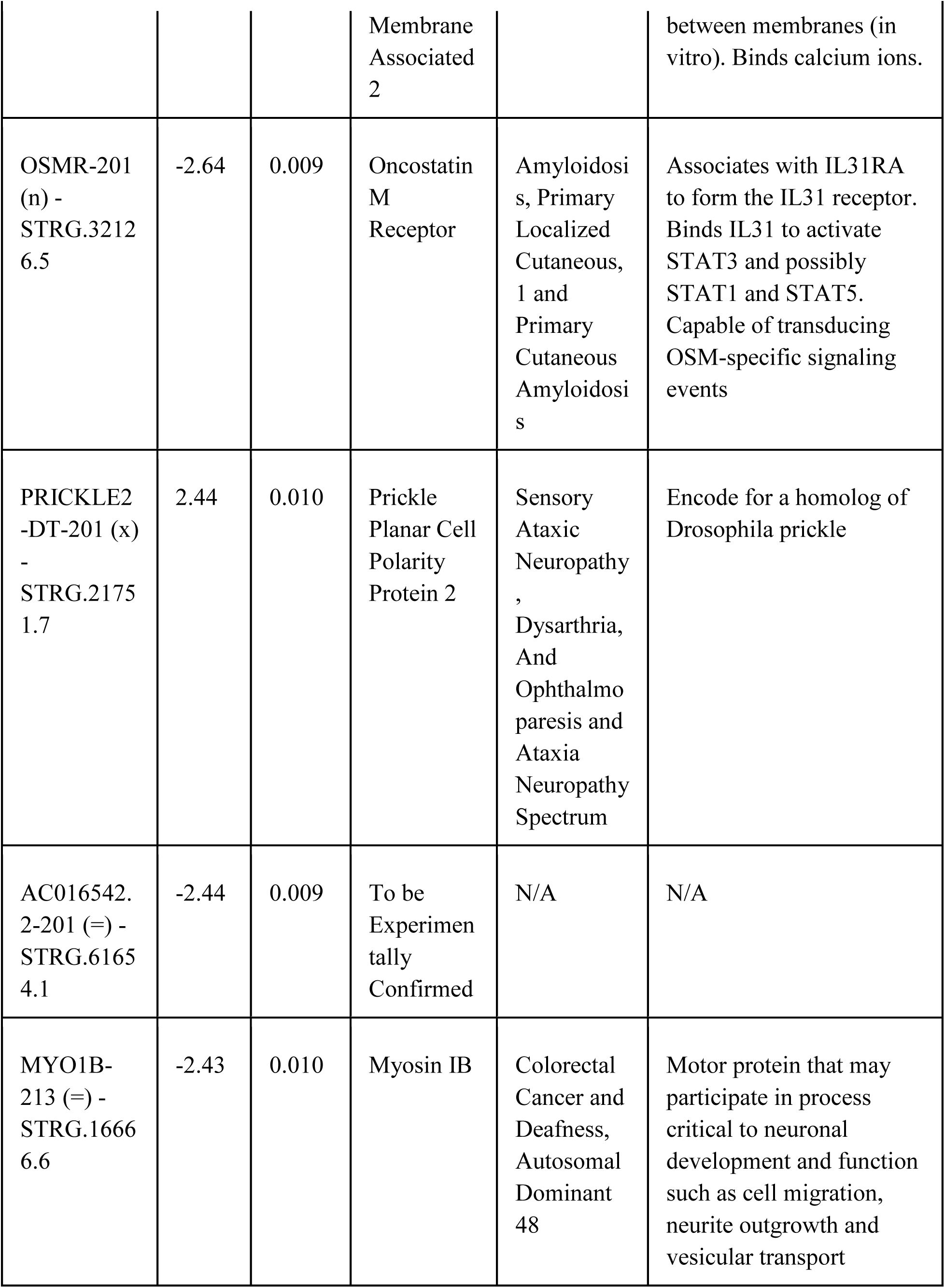
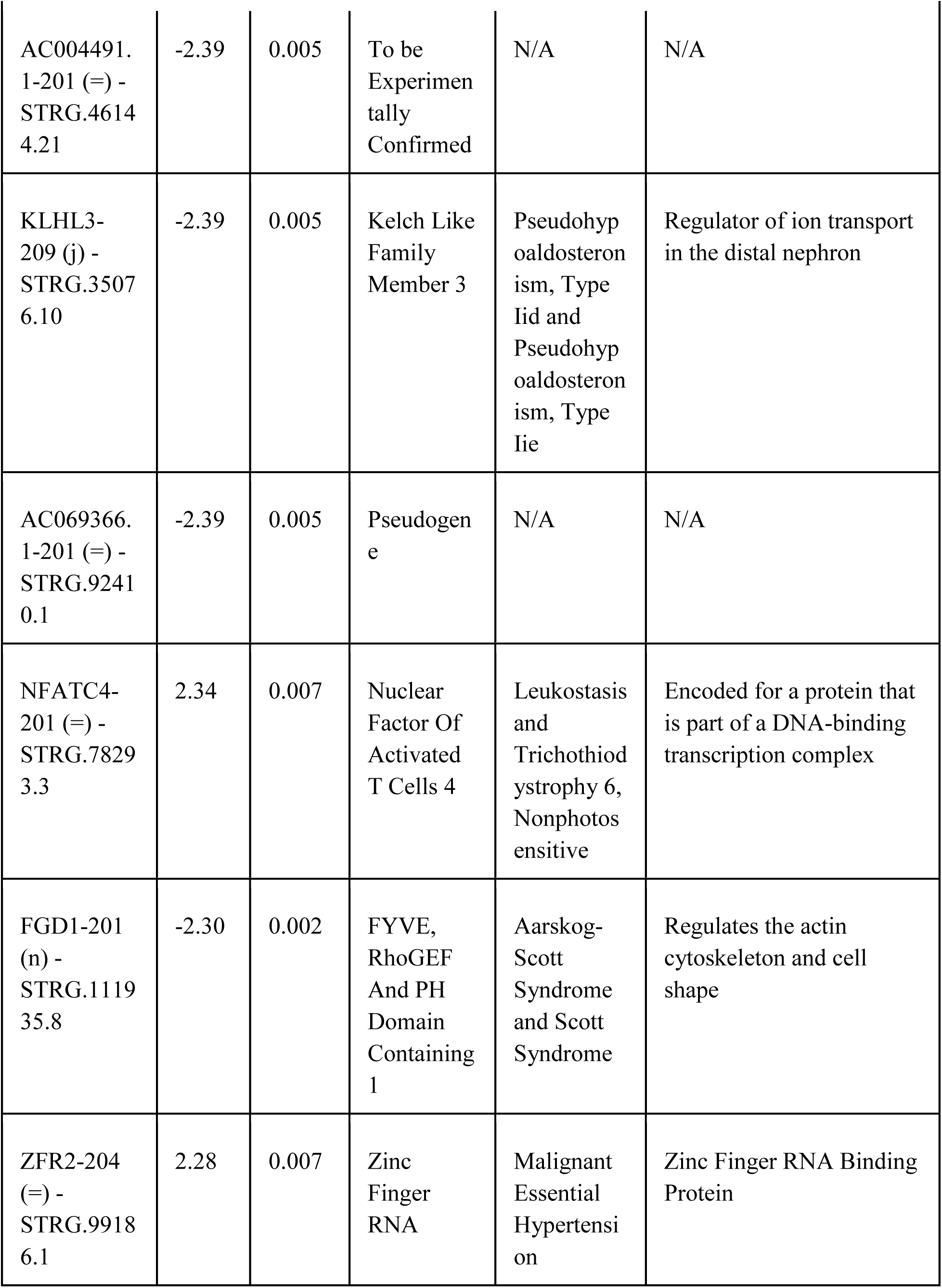
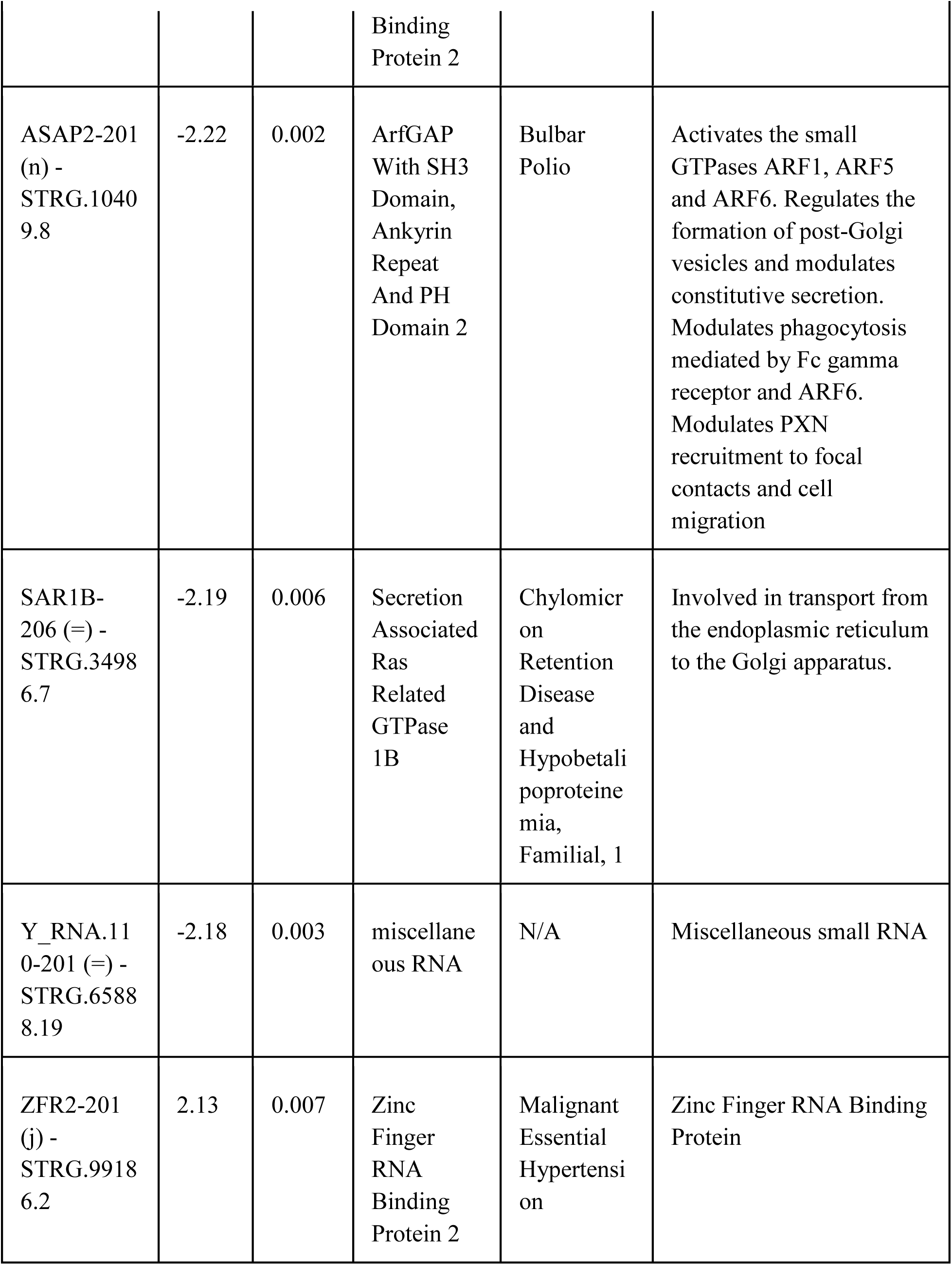
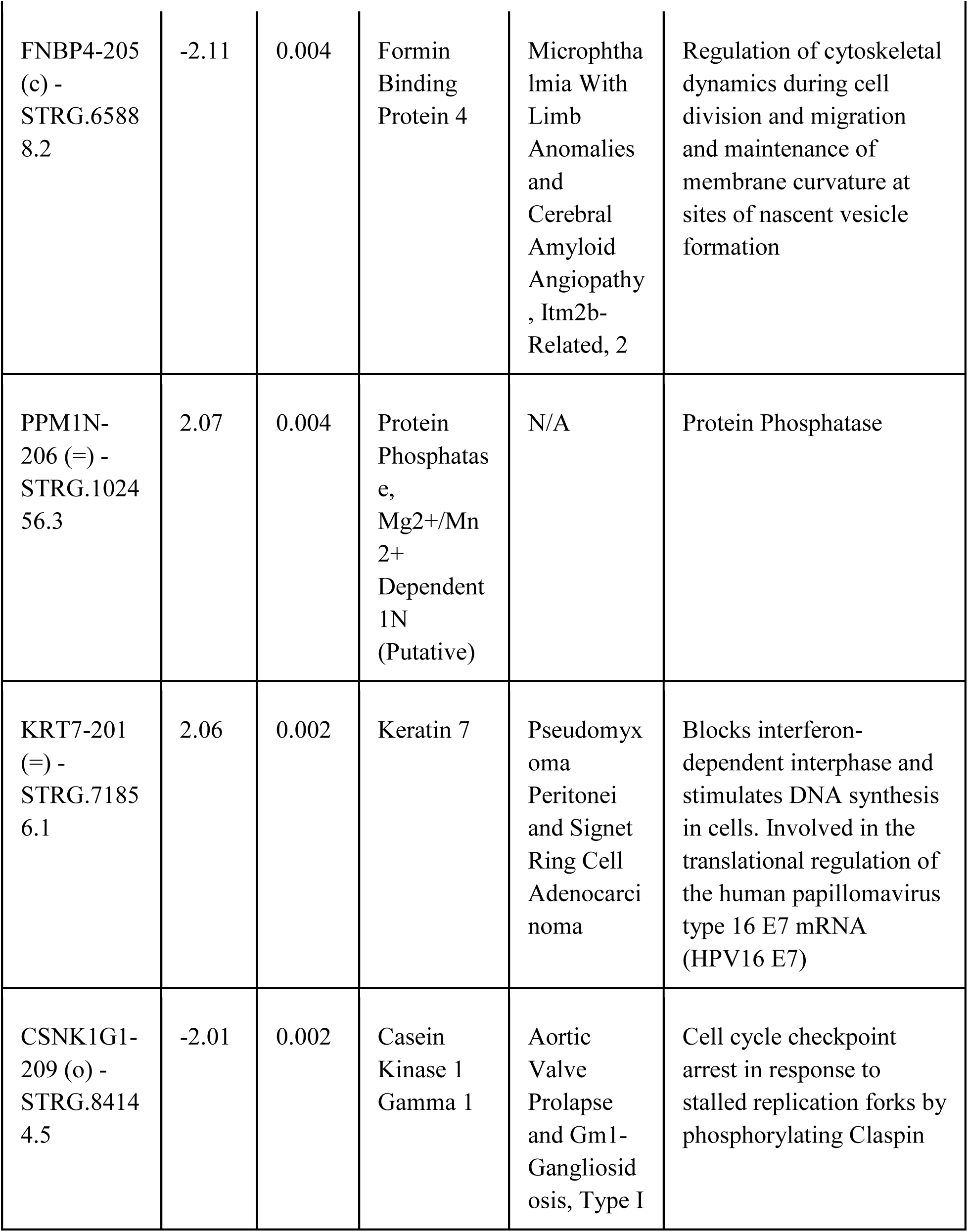
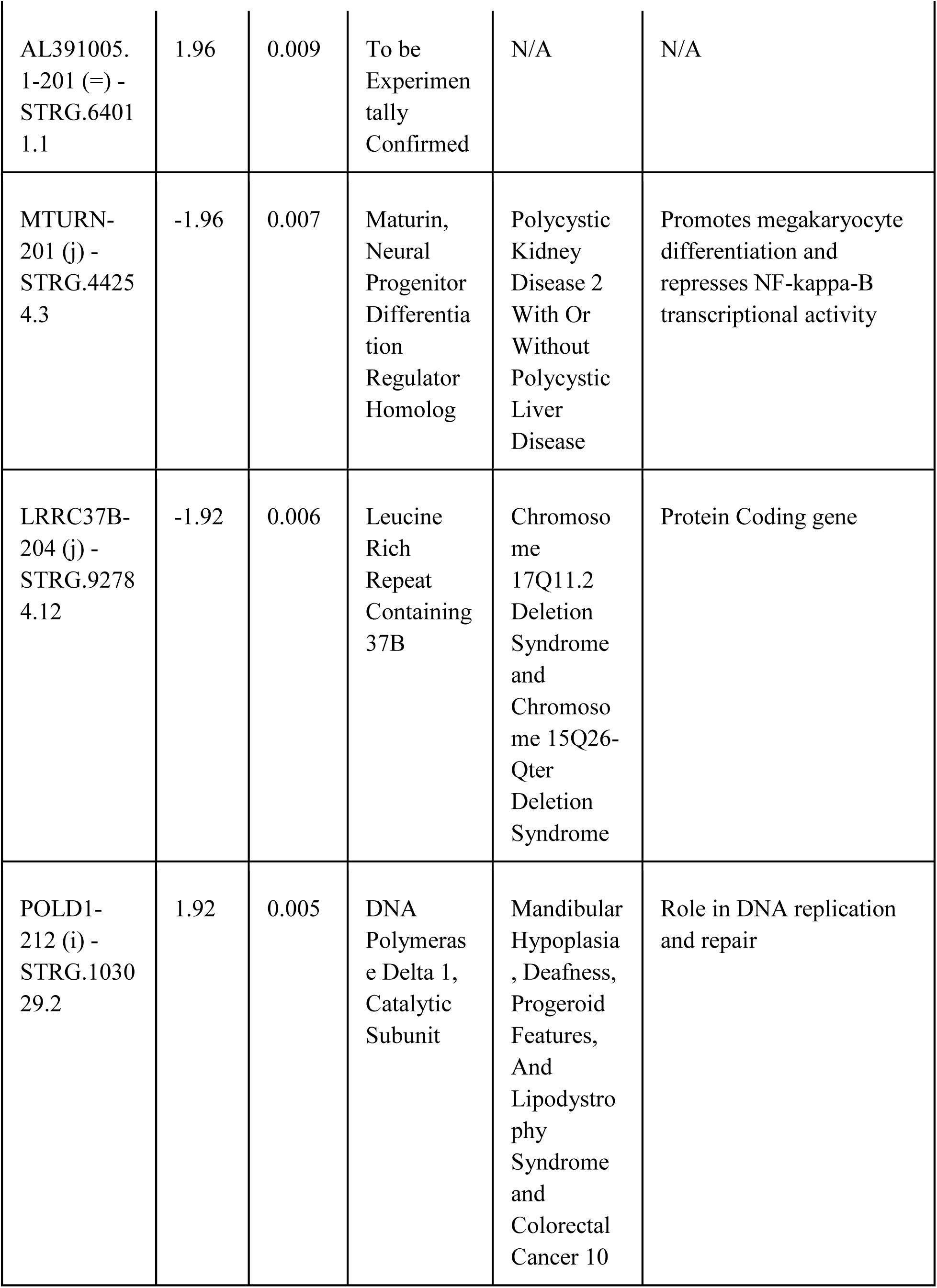
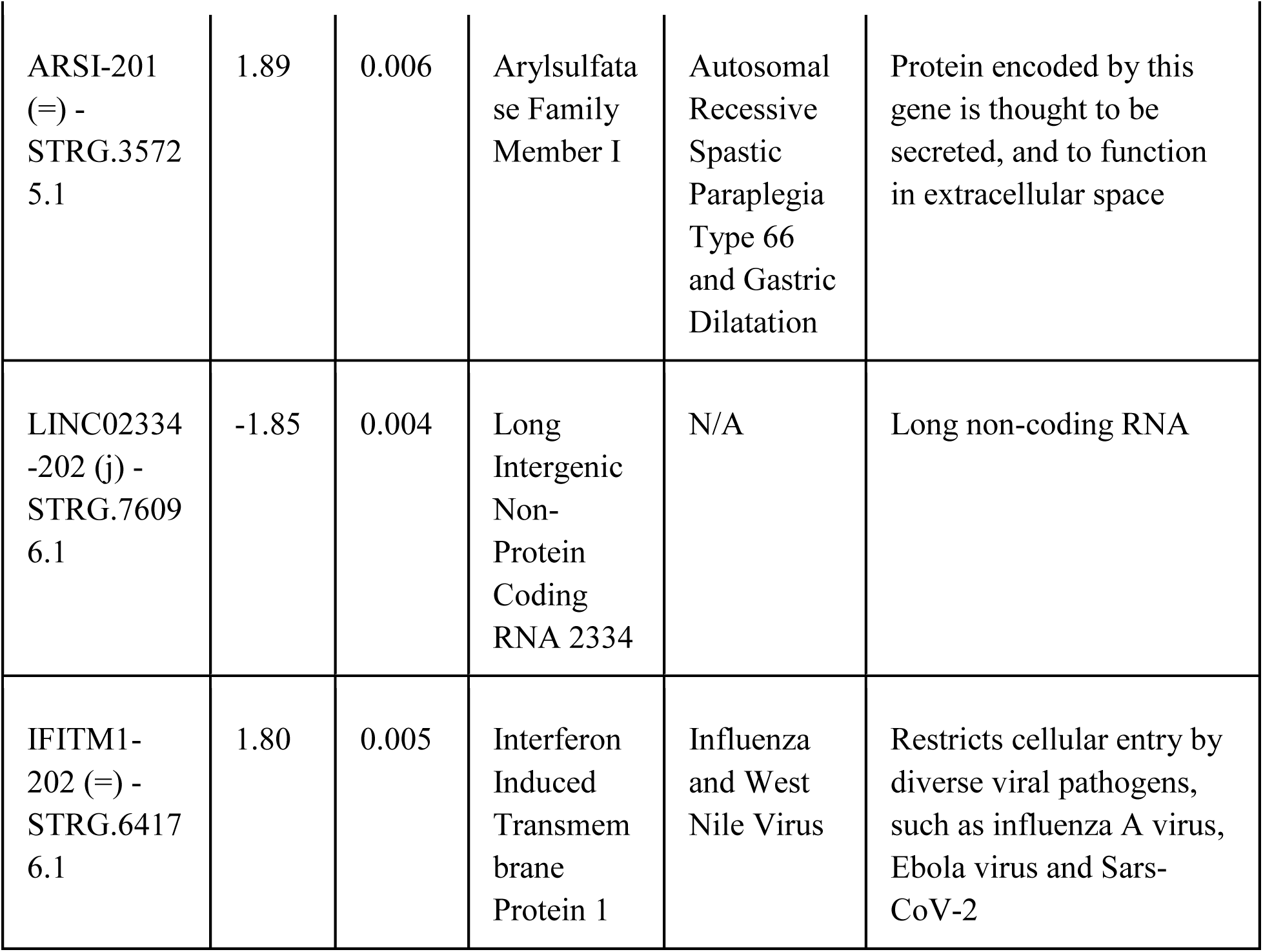
40 top differentially expressed transcripts in patient-derived anterior foregut cells. Long read single molecule cDNA sequencing analysis revealed a list of 173 transcripts differentially expressed in patient derived anterior foregut cells. The 40 most differentially expressed are described here (40 transcripts with the highest fold change value and batch-corrected P-value < 0.01).

**Supplementary Table 1b:**
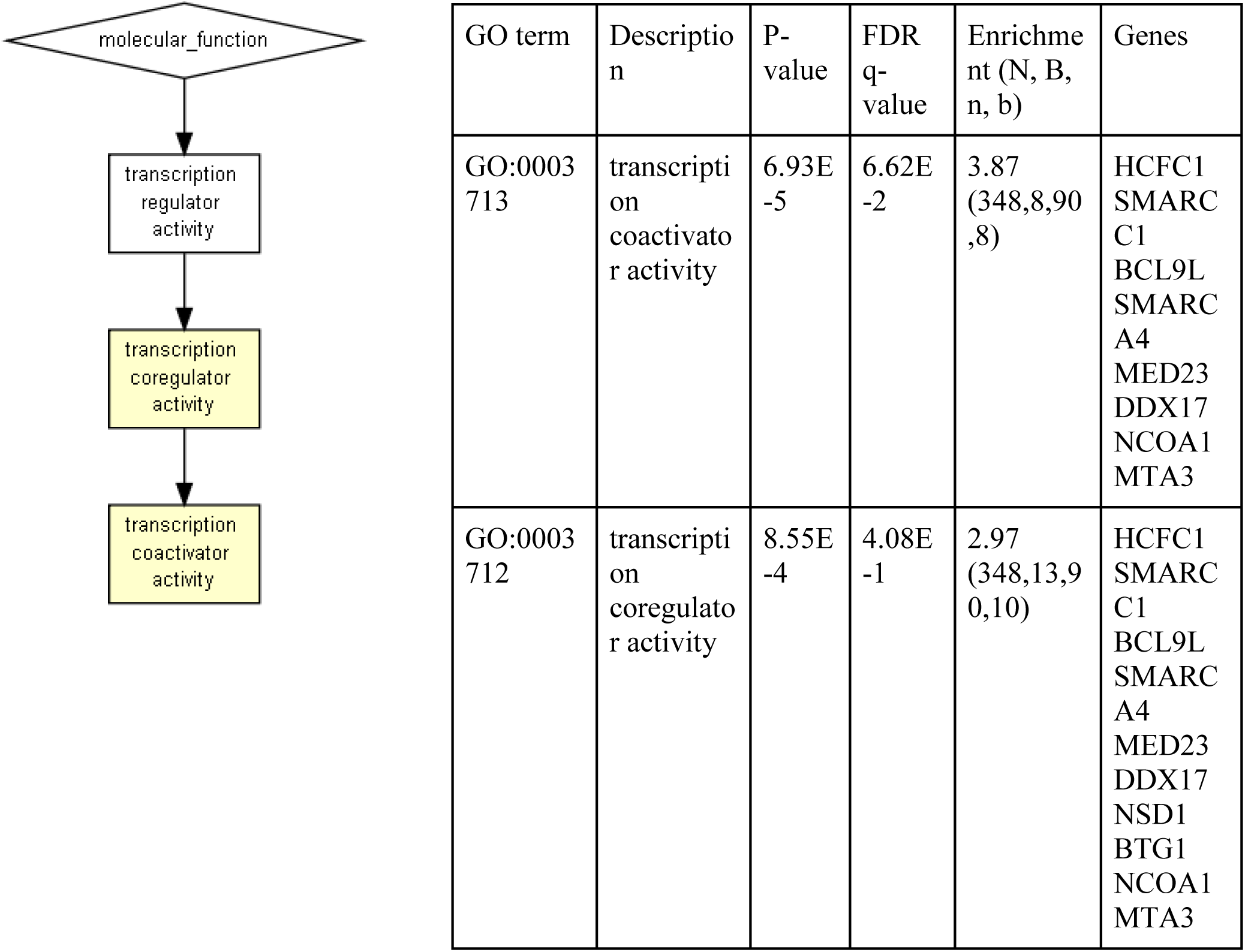
GO enrichment analysis results. Most differentially expressed transcripts uncovered by long read single molecule cDNA sequencing (Oxford Nanopore) revealed an enrichment of the transcription coactivator activity function with 8 genes involved in this function found differentially expressed (genes coding for transcripts with batch-corrected P-value < 0.01 and |log2(FoldChange)|>0.5 analyzed with GOrilla from Gene Ontology (Eden et al., 2009)).

| GO term | Description | P-value | FDR q-value | Enrichment (N, B, n, b) | Genes |
| --- | --- | --- | --- | --- | --- |
| GO:0003713 | transcription coactivator activity | 6.93E-5 | 6.62E-2 | 3.87 (348,8,90,8) | HCFC1<br>SMARCC1<br>BCL9L<br>SMARCA4<br>MED23<br>DDX17<br>NCOA1<br>MTA3 |
| GO:0003712 | transcription coregulator activity | 8.55E-4 | 4.08E-1 | 2.97 (348,13,90,10) | HCFC1<br>SMARCC1<br>BCL9L<br>SMARCA4<br>MED23<br>DDX17<br>NSD1<br>BTG1<br>NCOA1<br>MTA3 |

## References

Billmyre, K. K., M. Hutson, and J. Klingensmith. 2015. ’One shall become two: Separation of the esophagus and trachea from the common foregut tube’, Dev Dyn, 244: 277–88.

Brosens, E., J. F. Felix, A. Boerema-de Munck, E. M. de Jong, E. M. Lodder, S. Swagemakers, M. Buscop-van Kempen, R. R. de Krijger, R. M. H. Wijnen, IJcken W. F. J. van, P. van der Spek, A. de Klein, D. Tibboel, and R. J. Rottier. 2020. ’Histological, immunohistochemical and transcriptomic characterization of human tracheoesophageal fistulas’, PLoS One, 15: e0242167.

Brosens, E., M. Ploeg, Y. van Bever, A. E. Koopmans, I. Jsselstijn H, R. J. Rottier, R. Wijnen, D. Tibboel, and A. de Klein. 2014. ’Clinical and etiological heterogeneity in patients with tracheo-esophageal malformations and associated anomalies’, Eur J Med Genet, 57: 440–52.

Cho, S. G., Y. H. Lee, H. S. Park, K. Ryoo, K. W. Kang, J. Park, S. J. Eom, M. J. Kim, T. S. Chang, S. Y. Choi, J. Shim, Y. Kim, M. S. Dong, M. J. Lee, S. G. Kim, H. Ichijo, and E. J. Choi. 2001. ’Glutathione S-transferase mu modulates the stress-activated signals by suppressing apoptosis signal-regulating kinase 1’, J Biol Chem, 276: 12749–55.

Cho, S. H., I. Y. Kuo, P. F. Lu, H. T. Tzeng, W. W. Lai, W. C. Su, and Y. C. Wang. 2018. ’Rab37 mediates exocytosis of secreted frizzled-related protein 1 to inhibit Wnt signaling and thus suppress lung cancer stemness’, Cell Death Dis, 9: 868.

Clark, D. C. 1999. ’Esophageal atresia and tracheoesophageal fistula’, Am Fam Physician, 59: 910-6, 19–20.

DeWard, A. D., J. Cramer, and E. Lagasse. 2014. ’Cellular heterogeneity in the mouse esophagus implicates the presence of a nonquiescent epithelial stem cell population’, Cell Rep, 9: 701–11.

Domyan, E. T., E. Ferretti, K. Throckmorton, Y. Mishina, S. K. Nicolis, and X. Sun. 2011. ’Signaling through BMP receptors promotes respiratory identity in the foregut via repression of Sox2’, Development, 138: 971–81.

Eden, E., R. Navon, I. Steinfeld, D. Lipson, and Z. Yakhini. 2009. ’GOrilla: a tool for discovery and visualization of enriched GO terms in ranked gene lists’, BMC Bioinformatics, 10: 48.

Edwards, N. A., V. Shacham-Silverberg, L. Weitz, P. S. Kingma, Y. Shen, J. M. Wells, W. K. Chung, and A. M. Zorn. 2021. ’Developmental basis of trachea-esophageal birth defects’, Dev Biol, 477: 85–97.

Fausett, S. R., and J. Klingensmith. 2012. ’Compartmentalization of the foregut tube: developmental origins of the trachea and esophagus’, Wiley Interdiscip Rev Dev Biol, 1: 184–202.

Filonzi, L., C. Magnani, G. L. de’ Angelis, S. Dallaglio, and F. Nonnis Marzano. 2010. ’Evidence that polymorphic deletion of the glutathione S-transferase gene, GSTM1, is associated with esophageal atresia’, Birth Defects Res A Clin Mol Teratol, 88: 743–7.

Frankish, A., M. Diekhans, A. M. Ferreira, R. Johnson, I. Jungreis, J. Loveland, J. M. Mudge, C. Sisu, J. Wright, J. Armstrong, I. Barnes, A. Berry, A. Bignell, S. Carbonell Sala, J. Chrast, F. Cunningham, T. Di Domenico, S. Donaldson, I. T. Fiddes, C. Garcia Giron, J. M. Gonzalez, T. Grego, M. Hardy, T. Hourlier, T. Hunt, O. G. Izuogu, J. Lagarde, F. J. Martin, L. Martinez, S. Mohanan, P. Muir, F. C. P. Navarro, A. Parker, B. Pei, F. Pozo, M. Ruffier, B. M. Schmitt, E. Stapleton, M. M. Suner, I. Sycheva, B. Uszczynska-Ratajczak, J. Xu, A. Yates, D. Zerbino, Y. Zhang, B. Aken, J. S. Choudhary, M. Gerstein, R. Guigo, T. J. P. Hubbard, M. Kellis, B. Paten, A. Reymond, M. L. Tress, and P. Flicek. 2019. ’GENCODE reference annotation for the human and mouse genomes’, Nucleic Acids Res, 47: D766–D73.

Giroux, V., A. A. Lento, M. Islam, J. R. Pitarresi, A. Kharbanda, K. E. Hamilton, K. A. Whelan, A. Long, B. Rhoades, Q. Tang, H. Nakagawa, C. J. Lengner, A. J. Bass, E. P. Wileyto, A. J. Klein-Szanto, T. C. Wang, and A. K. Rustgi. 2017. ’Long-lived keratin 15+ esophageal progenitor cells contribute to homeostasis and regeneration’, J Clin Invest, 127: 2378–91.

Guyot, B., and V. Maguer-Satta. 2016. ’Blocking TGF-beta and BMP SMAD-dependent cell differentiation is a master key to expand all kinds of epithelial stem cells’, Stem Cell Investig, 3: 88.

Han, L., P. Chaturvedi, K. Kishimoto, H. Koike, T. Nasr, K. Iwasawa, K. Giesbrecht, P. C. Witcher, A. Eicher, L. Haines, Y. Lee, J. M. Shannon, M. Morimoto, J. M. Wells, T. Takebe, and A. M. Zorn. 2020. ’Single cell transcriptomics identifies a signaling network coordinating endoderm and mesoderm diversification during foregut organogenesis’, Nat Commun, 11: 4158.

Hrstka, S. C., X. Li, T. J. Nelson, and Group Wanek Program Genetics Pipeline. 2017. ’NOTCH1-Dependent Nitric Oxide Signaling Deficiency in Hypoplastic Left Heart Syndrome Revealed Through Patient-Specific Phenotypes Detected in Bioengineered Cardiogenesis’, Stem Cells, 35: 1106–19.

Huang, S. X., M. D. Green, A. T. de Carvalho, M. Mumau, Y. W. Chen, S. L. D’Souza, and H. W. Snoeck. 2015. ’The in vitro generation of lung and airway progenitor cells from human pluripotent stem cells’, Nat Protoc, 10: 413–25.

Huang, S. X., M. N. Islam, J. O’Neill, Z. Hu, Y. G. Yang, Y. W. Chen, M. Mumau, M. D. Green, G. Vunjak-Novakovic, J. Bhattacharya, and H. W. Snoeck. 2014. ’Efficient generation of lung and airway epithelial cells from human pluripotent stem cells’, Nat Biotechnol, 32: 84–91.

Ioannides, A. S., V. Massa, E. Ferraro, F. Cecconi, L. Spitz, D. J. Henderson, and A. J. Copp. 2010. ’Foregut separation and tracheo-oesophageal malformations: the role of tracheal outgrowth, dorso-ventral patterning and programmed cell death’, Dev Biol, 337: 351–62.

Karagiannis, P., K. Takahashi, M. Saito, Y. Yoshida, K. Okita, A. Watanabe, H. Inoue, J. K. Yamashita, M. Todani, M. Nakagawa, M. Osawa, Y. Yashiro, S. Yamanaka, and K. Osafune. 2019. ’Induced Pluripotent Stem Cells and Their Use in Human Models of Disease and Development’, Physiol Rev, 99: 79–114.

Kim, E., M. Jiang, H. Huang, Y. Zhang, N. Tjota, X. Gao, J. Robert, N. Gilmore, L. Gan, and J. Que. 2019. ’Isl1 Regulation of Nkx2.1 in the Early Foregut Epithelium Is Required for Trachea-Esophageal Separation and Lung Lobation’, Dev Cell, 51: 675–83 e4.

Kovaka, S., A. V. Zimin, G. M. Pertea, R. Razaghi, S. L. Salzberg, and M. Pertea. 2019. ’Transcriptome assembly from long-read RNA-seq alignments with StringTie2’, Genome Biol, 20: 278.

Kuwahara, A., A. E. Lewis, C. Coombes, F. S. Leung, M. Percharde, and J. O. Bush. 2020. ’Delineating the early transcriptional specification of the mammalian trachea and esophagus’, Elife, 9.

Leek, J.T., Johnson, W.E., Parker, H.S., Fertig, E.J., Jaffe, A.E., Zhang, Y., Storey, J.D., Torres, L.C. 2021. ’sva: Surrogate Variable Analysis.’, R package version 3.42.0.

Li, H. 2018. ’Minimap2: pairwise alignment for nucleotide sequences’, Bioinformatics, 34: 3094–100.

Li, H., B. Handsaker, A. Wysoker, T. Fennell, J. Ruan, N. Homer, G. Marth, G. Abecasis, R. Durbin, and Subgroup Genome Project Data Processing. 2009. ’The Sequence Alignment/Map format and SAMtools’, Bioinformatics, 25: 2078–9.

Li, Y., Y. Litingtung, P. Ten Dijke, and C. Chiang. 2007. ’Aberrant Bmp signaling and notochord delamination in the pathogenesis of esophageal atresia’, Dev Dyn, 236: 746–54.

Love, M. I., W. Huber, and S. Anders. 2014. ’Moderated estimation of fold change and dispersion for RNA-seq data with DESeq2’, Genome Biol, 15: 550.

Matsuno, K., S. I. Mae, C. Okada, M. Nakamura, A. Watanabe, T. Toyoda, E. Uchida, and K. Osafune. 2016. ’Redefining definitive endoderm subtypes by robust induction of human induced pluripotent stem cells’, Differentiation, 92: 281–90.

Miao, Y., L. Tian, M. Martin, S. L. Paige, F. X. Galdos, J. Li, A. Klein, H. Zhang, N. Ma, Y. Wei, M. Stewart, S. Lee, J. R. Moonen, B. Zhang, P. Grossfeld, S. Mital, D. Chitayat, J. C. Wu, M. Rabinovitch, T. J. Nelson, S. Nie, S. M. Wu, and M. Gu. 2020. ’Intrinsic Endocardial Defects Contribute to Hypoplastic Left Heart Syndrome’, Cell Stem Cell, 27: 574–89.e8.

Minoo, P., G. Su, H. Drum, P. Bringas, and S. Kimura. 1999. ’Defects in tracheoesophageal and lung morphogenesis in Nkx2.1(-/-) mouse embryos’, Dev Biol, 209: 60–71.

Nam, K. T., H. J. Lee, J. J. Smith, L. A. Lapierre, V. P. Kamath, X. Chen, B. J. Aronow, T. J. Yeatman, S. G. Bhartur, B. C. Calhoun, B. Condie, N. R. Manley, R. D. Beauchamp, R. J. Coffey, and J. R. Goldenring. 2010. ’Loss of Rab25 promotes the development of intestinal neoplasia in mice and is associated with human colorectal adenocarcinomas’, J Clin Invest, 120: 840–9.

Nasr, T., P. Mancini, S. A. Rankin, N. A. Edwards, Z. N. Agricola, A. P. Kenny, J. L. Kinney, K. Daniels, J. Vardanyan, L. Han, S. L. Trisno, S. W. Cha, J. M. Wells, M. J. Kofron, and A. M. Zorn. 2019. ’Endosome-Mediated Epithelial Remodeling Downstream of Hedgehog-Gli Is Required for Tracheoesophageal Separation’, Dev Cell, 51: 665–74 e6.

Patro, R., G. Duggal, M. I. Love, R. A. Irizarry, and C. Kingsford. 2017. ’Salmon provides fast and bias-aware quantification of transcript expression’, Nat Methods, 14: 417–19.

Pertea, G., and M. Pertea. 2020. ’GFF Utilities: GffRead and GffCompare’, F1000Res, 9.

Que, J., M. Choi, J. W. Ziel, J. Klingensmith, and B. L. Hogan. 2006. ’Morphogenesis of the trachea and esophagus: current players and new roles for noggin and Bmps’, Differentiation, 74: 422–37.

Que, J., T. Okubo, J. R. Goldenring, K. T. Nam, R. Kurotani, E. E. Morrisey, O. Taranova, L. H. Pevny, and B. L. Hogan. 2007. ’Multiple dose-dependent roles for Sox2 in the patterning and differentiation of anterior foregut endoderm’, Development, 134: 2521–31.

R Core Team. 2021. ’R: A language and environment for statistical computing.’ R Foundation for Statistical Computing, Vienna, Austria.

Raad, S., A. David, J. Que, and C. Faure. 2020. ’Genetic Mouse Models and Induced Pluripotent Stem Cells for Studying Tracheal-Esophageal Separation and Esophageal Development’, Stem Cells Dev, 29: 953–66.

Rosekrans, S. L., B. Baan, V. Muncan, and G. R. van den Brink. 2015. ’Esophageal development and epithelial homeostasis’, Am J Physiol Gastrointest Liver Physiol, 309: G216–28.

Rowe, R. G., and G. Q. Daley. 2019. ’Induced pluripotent stem cells in disease modelling and drug discovery’, Nat Rev Genet, 20: 377–88.

Shahryari, A., M. R. Rafiee, Y. Fouani, N. A. Oliae, N. M. Samaei, M. Shafiee, S. Semnani, M. Vasei, and S. J. Mowla. 2014. ’Two novel splice variants of SOX2OT, SOX2OT-S1, and SOX2OT-S2 are coupregulated with SOX2 and OCT4 in esophageal squamous cell carcinoma’, Stem Cells, 32: 126–34.

Shelton, M., A. Kocharyan, J. Liu, I. S. Skerjanc, and W. L. Stanford. 2016. ’Robust generation and expansion of skeletal muscle progenitors and myocytes from human pluripotent stem cells’, Methods, 101: 73–84.

Stoll, C., Y. Alembik, B. Dott, and M. P. Roth. 2009. ’Associated malformations in patients with esophageal atresia’, Eur J Med Genet, 52: 287–90.

Teramoto, M., R. Sugawara, K. Minegishi, M. Uchikawa, T. Takemoto, A. Kuroiwa, Y. Ishii, and H. Kondoh. 2020. ’The absence of SOX2 in the anterior foregut alters the esophagus into trachea and bronchi in both epithelial and mesenchymal components’, Biol Open, 9: 048728.

Trisno, S. L., K. E. D. Philo, K. W. McCracken, E. M. Cata, S. Ruiz-Torres, S. A. Rankin, L. Han, T. Nasr, P. Chaturvedi, M. E. Rothenberg, M. A. Mandegar, S. I. Wells, A. M. Zorn, and J. M. Wells. 2018. ’Esophageal Organoids from Human Pluripotent Stem Cells Delineate Sox2 Functions during Esophageal Specification’, Cell Stem Cell, 23: 501–15 e7.

van Lennep, M., M. M. J. Singendonk, L. Dall’Oglio, F. Gottrand, U. Krishnan, S. W. J. Terheggen-Lagro, T. I. Omari, M. A. Benninga, and M. P. van Wijk. 2019. ’Oesophageal atresia’, Nat Rev Dis Primers, 5: 26.

Wang, J., P. R. Ahimaz, S. Hashemifar, J. Khlevner, J. A. Picoraro, W. Middlesworth, M. M. Elfiky, J. Que, Y. Shen, and W. K. Chung. 2021. ’Novel candidate genes in esophageal atresia/tracheoesophageal fistula identified by exome sequencing’, Eur J Hum Genet, 29: 122–30.

Wells, J. M., and D. A. Melton. 1999. ’Vertebrate endoderm development’, Annu Rev Cell Dev Biol, 15: 393–410.

Woo, J., I. Miletich, B. M. Kim, P. T. Sharpe, and R. A. Shivdasani. 2011. ’Barx1-mediated inhibition of Wnt signaling in the mouse thoracic foregut controls tracheo-esophageal septation and epithelial differentiation’, PLoS One, 6: e22493.

Yang, C., Y. Xu, M. Yu, D. Lee, S. Alharti, N. Hellen, N. Ahmad Shaik, B. Banaganapalli, H. Sheikh Ali Mohamoud, R. Elango, S. Przyborski, G. Tenin, S. Williams, J. O’Sullivan, O. O. Al-Radi, J. Atta, S. E. Harding, B. Keavney, M. Lako, and L. Armstrong. 2017. ’Induced pluripotent stem cell modelling of HLHS underlines the contribution of dysfunctional NOTCH signalling to impaired cardiogenesis’, Hum Mol Genet, 26: 3031–45.

Zhang, Y., Y. Yang, M. Jiang, S. X. Huang, W. Zhang, D. Al Alam, S. Danopoulos, M. Mori, Y. W. Chen, R. Balasubramanian, S. M. Chuva de Sousa Lopes, C. Serra, M. Bialecka, E. Kim, S. Lin, A. L. R. Toste de Carvalho, P. N. Riccio, W. V. Cardoso, X. Zhang, H. W. Snoeck, and J. Que. 2018. ’3D Modeling of Esophageal Development using Human PSC-Derived Basal Progenitors Reveals a Critical Role for Notch Signaling’, Cell Stem Cell, 23: 516–29 e5.

